# AlphaGEM Enables Precise Genome-Scale Metabolic Modelling by Integrating Protein Structure Alignment with deep-learning-based Dark Metabolism Mining

**DOI:** 10.1101/2025.07.21.665674

**Authors:** Weishang Han, Luchi Xiao, Haocheng Sun, Guangming Xiang, Qianxi Jia, Haoyu Wang, Boyang Ji, Cheng Zhang, Eduard J. Kerkhoven, Jens Nielsen, Hongzhong Lu

## Abstract

Constructing high-quality genome-scale metabolic models (GEMs) for less-studied species remains challenging. To address this, we developed AlphaGEM, a versatile toolbox leveraging proteome-scale structural alignment and deep-learning-based predictions for efficient genomic mining to generate GEMs ready for applications. Our findings show that the structural alignment or protein-language-model-based prediction (i.e., PLMSearch), could identify more homologous protein relationships than sequence-blast-based alignment, contributing to the accurate profiling of metabolism from target organisms. Additionally, AlphaGEM encompasses an ensemble procedure empowered by multiple deep learning toolboxes to effectively mine the dark metabolic functions encoded by nonhomologous proteins, significantly expanding species-specific metabolic networks. We demonstrate AlphaGEM’s accuracy by building GEMs for eukaryotes (e.g., *S. pombe*, *C. albicans*) and prokaryotes (e.g., *K. pneumoniae*, *B. subtilis*), achieving predictions comparable to manually curated models while outperforming existing tools. AlphaGEM also successfully reconstructs GEMs for *M. musculus* and *C. griseus*, showcasing its great potential for uncovering dark metabolism in complex mammals. Lastly, we demonstrate that AlphaGEM could facilitate the automatic GEMs reconstruction for 332 distinct yeast species with high prediction fidelity. In conclusion, AlphaGEM provides unprecedented opportunities for the precise, rapid construction of GEMs across diverse domains, which sets a solid foundation for universal functional analysis of non-model organisms having genome sequences available.

## Introduction

Genome-scale metabolic models (GEMs) are powerful computational frameworks for analyzing cellular metabolism, allowing them to predict phenotypic outcomes under genetic and environmental perturbations^1–4^. GEMs have a variety of applications, including metabolic engineering for optimizing microbial strains for biochemical production and drug discovery for precision medicine^4^, where GEMs help identify potential therapeutic targets by simulating cellular responses to perturbations^5–7^. Furthermore, GEMs allow for the investigation of metabolic interactions within microbial communities, providing insights into ecological dynamic evolution for the better adaptability^8^. Combining omics data with advanced algorithms further improves GEM predictive capabilities, which advances the studies in personalized medicine and synthetic biology^9–12^.

Currently, the increasing accumulation of large-scale genomic data from high-throughput sequencing experiment is in great demand of high-quality GEM for computational analysis of cellular metabolism^13^. Note that various GEMs for model organisms that are widely used have been going through a combination of semi-automated and manual curation over many years (i.e., yeast-GEM^14^, iML1515^15^, human-GEM^16^), where the quality of the model is improved with each iteration. However, the labour-intensive approach used for model organisms is not feasible to be applied for large-scale newly sequenced species. Meanwhile, the complexity and variety in metabolic networks across species seriously hinder the reconstruction of high-quality, organism-specific GEMs for many less characterized species^17^. Therefore, the development of powerful computational toolboxes capable of producing ready-to-use GEMs is of great value.

Until now, several automatic modeling tools, including CarveMe^18^, ModelSEED^10^, gapseq^19^, MERLIN^20^, Metadraft^21^ and AuReMe^22^ have been developed to facilitate the reconstruction of GEMs. However, these tools often produce models with suboptimal quality and accuracy, requiring manual corrections before ready usage^23^. CarveMe, for example, generates metabolic models for the target organism by manually building universal metabolic network templates^18^. Although the metabolic models it generates have more reactions without gene annotations and can be used directly for flux balance analysis (FBA), those GEMs frequently have low specificity, with high similarities between models from different species. Other template-based tools, including Metadraft and AuReMe, build models from predefined template models. These tools have the same issues as CarveMe, and they perform poorly when annotating proteins that do not have homologous proteins in the template model’s species, which possibly not reflecting species-specific metabolic characteristics. ModelSeed, MERLIN, and gapseq, on the other hand, begin with the genome, then utilize various bioinformatics tools to annotate genes, potentially improving the completeness of metabolic models. However, these toolboxes might be accompanied with poor protein annotation, which, to some extent, lowering the model quality^23^. Addressing these issues is highly critical to promote the usability of GEMs by the wider scientific community.

To increase the quality of GEMs and accurately characterize the metabolic diversity of numerous newly sequenced species, we created AlphaGEM, a versatile computational toolbox that integrating protein homology inference and dark metabolism mining to address the bottleneck existing in current toolboxes for automatic GEMs reconstruction. The performance of AlphaGEM was thoroughly assessed, and multiple case studies were used to demonstrate its advantages in creating GEMs for various organisms from the simple bacteria organism to the complex mammal organism. Overall, AlphaGEM enables high-quality and high-throughput GEM reconstruction by refining the protein function annotation leveraging structure and sequence alignment on top of a series of advanced deep learning models, which will certainly accelerate model-based simulation and analysis for a wide range of organisms.

## Results

### Overview of AlphaGEM framework

Recent advancements in deep learning offer unprecedented opportunities to determine gene and protein function with high resolution. We introduce AlphaGEM, a versatile framework for reconstructing high-quality GEMs for sequenced organisms through protein function inference derived from protein 3D structure comparison and protein language model prediction. Different from previous studies^10, 18^, which select mature universal models to rebuilt GEMs for non-model organisms, we utilize curated high-quality GEMs from model organisms as references— employing iML1515^15^ as the template model for prokaryotes, Yeast9^14^ for unicellular eukaryotes and fungi, and Human1^16^ for mammalian cells (Fig.1). Within AlphaGEM’s initial module, draft-GEMs could be automatically generated from the aforementioned reference models based on the inferred protein homologous relationships between the model and non-model organisms. Departing from conventional approaches that depend solely on sequence similarity to establish homology, AlphaGEM implements two independent strategies: (1) using genome-scale homologous relationships analysis via sequence similarity and protein structure similarity when proteome-scale 3D structures are available, and (2) leveraging pre-training protein language models, i.e., PLMSearch^24^, to infer homologous relationships when proteome-scale structures are unavailable (Fig. 1, Methods), both will be detailed respectively in the following sections.

**Figure 1.**
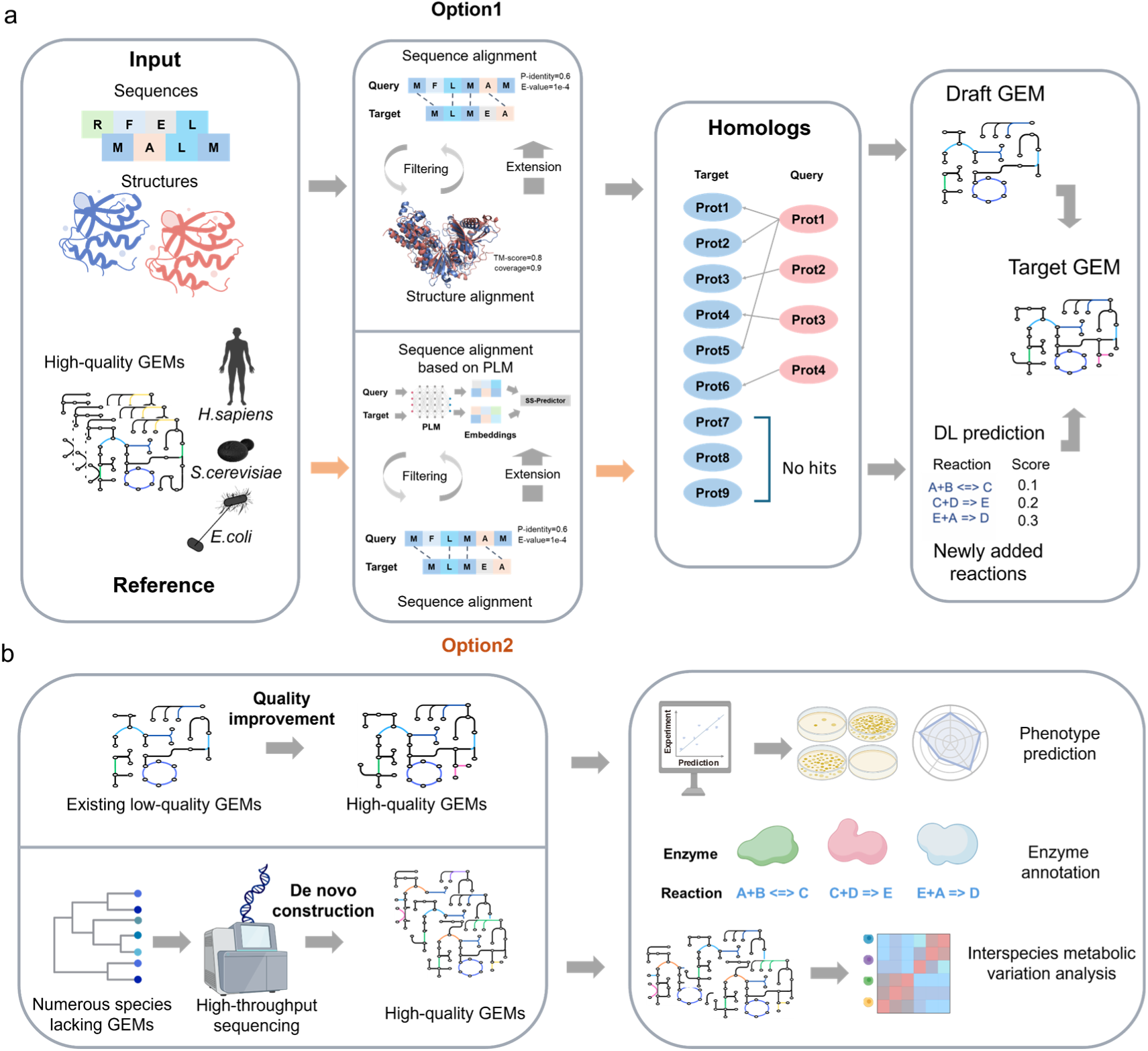
Framework used in AlphaGEM for constructing high-quality GEMs for non-model organisms. With model organisms’ high-quality GEMs as references, AlphaGEM integrates biological sequence and protein structural information to identify homologous relationships via combined sequence/structure alignment (or PLMSearch when structure is unavailable). This homologous information drives the construction of a draft GEM. Subsequently, nonhomologous proteins undergo functional annotation, scoring, and filtering using multiple deep learning tools, and are integrated into the draft GEM. Finally, the above outputs are combined to obtain ready-to-use GEMs for specific organisms (a). Typical applications of AlphaGEM. For example, AlphaGEM enables optimization of low-quality GEMs or de novo reconstruction of GEMs for non-model organisms. GEMs generated by AlphaGEM could be used to perform systematic, quantitative, and predictive analyses of cellular metabolism, mine the enzyme function and probe the interaction of distinct strains within a community (b).

Although reference models were utilized to generate draft GEMs, there still exist amounts of species-specific metabolic reactions and pathways in the target organisms. To enhance metabolic coverage, we implemented a bottom-up strategy by annotating nonhomologous proteins from target organisms. Traditional sequence-based annotation tools often exhibit limited accuracy^25^, whereas recent advances in deep learning have provided more robust and scalable solutions. Here, we developed an ensemble pipeline with aid of several advanced computational tools, such as CLEAN^26^, DeepECtransformer^27^ and eggNOG-mapper^28^, to annotate nonhomologous proteins encoding metabolic proteins within the genomes from the input organisms (Fig. 1), thereby significantly expanding the scope of GEMs for non-model organisms. Lastly, to address the potential gaps in metabolic network resulting from inaccuracies existing in sequencing or assembly, we performed gap-filling with reference models as templates using iterative gap-filling and evaluation^17^. Collectively, AlphaGEM seamlessly integrates a suite of advanced modules to provide a reliable and effective solution for modeling the metabolism of newly sequenced organisms.

### Cross-species homologous relationships inference by integrating sequence and structure information with AlphaGEM

The quality of initial draft-GEMs critically depends on the reliable annotation of protein homologous relationships between input and reference proteins. Traditional approaches typically rely on sequence alignment tools like HMMER^29^, blastp^30^, and OrthoFinder^31^. However, using sequence alignment to establish protein homologous relationships between distinct species is frequently insufficient, as certain proteins with similar functions may not be identified due to limited sequence similarity^32, 33^. To address this limitation, we incorporated proteome-scale structural alignment to refine homologous relationships inference beyond what is possible with sequence alignment alone (Methods, Supplemental Fig. 1a).

To validate the reliability of the homologous relationships obtained through the above procedures, we evaluated the performance of AlphaGEM in inference of homologous relationships using proteomes from two representative fungal species, *Candida albicans* and *Saccharomyces cerevisiae*, the latter of which was set the reference organism. At the first glance, we observed that protein pairs exhibiting high sequence similarity generally align with those showing high structural similarity (Fig. 2a). However, a significant number of protein pairs with high structural similarity have relatively low sequence similarity (Fig. 2b). Therefore, focusing on GEMs, we compared the homologous groups of metabolic proteins inferred by different steps encompassed within AlphaGEM: 1) OrthoFinder based grouping, 2) US-align based filtering, and 3) Foldseek-based expansion (Methods, Fig. 2c). Compared with the first step, US-align based structural clustering approach could successfully decrease false homologs from 63 to 11 and improve the accuracy from 94.91% to 98.88% (Supplemental Fig. 2c, 2f). It shows that structure-based comparison could help to identify additional homologous relationships as some of homologous pairs are more conserved in protein 3D structure than that in protein sequence (Fig. 2d-2e). Importantly, it is found that sequence-based homologous protein pairs with high percent identity (identity) but low TM-score could have distinct metabolic functions (Fig. 2f-2g), showcasing the necessity of filtering out low-quality orthologous protein relationships using structural alignment. Overall, our whole pipeline encompassing the above three steps identified more homologous pairs (an increase by 26.35%) than the sequence-based clustering approach due to the extra Foldseek-based structure comparison at proteome scale. However, we found the increased homologous pairs could result in a slight decrease in accuracy from 94.91% to 94.71% (Fig. 2c).

**Figure 2.**
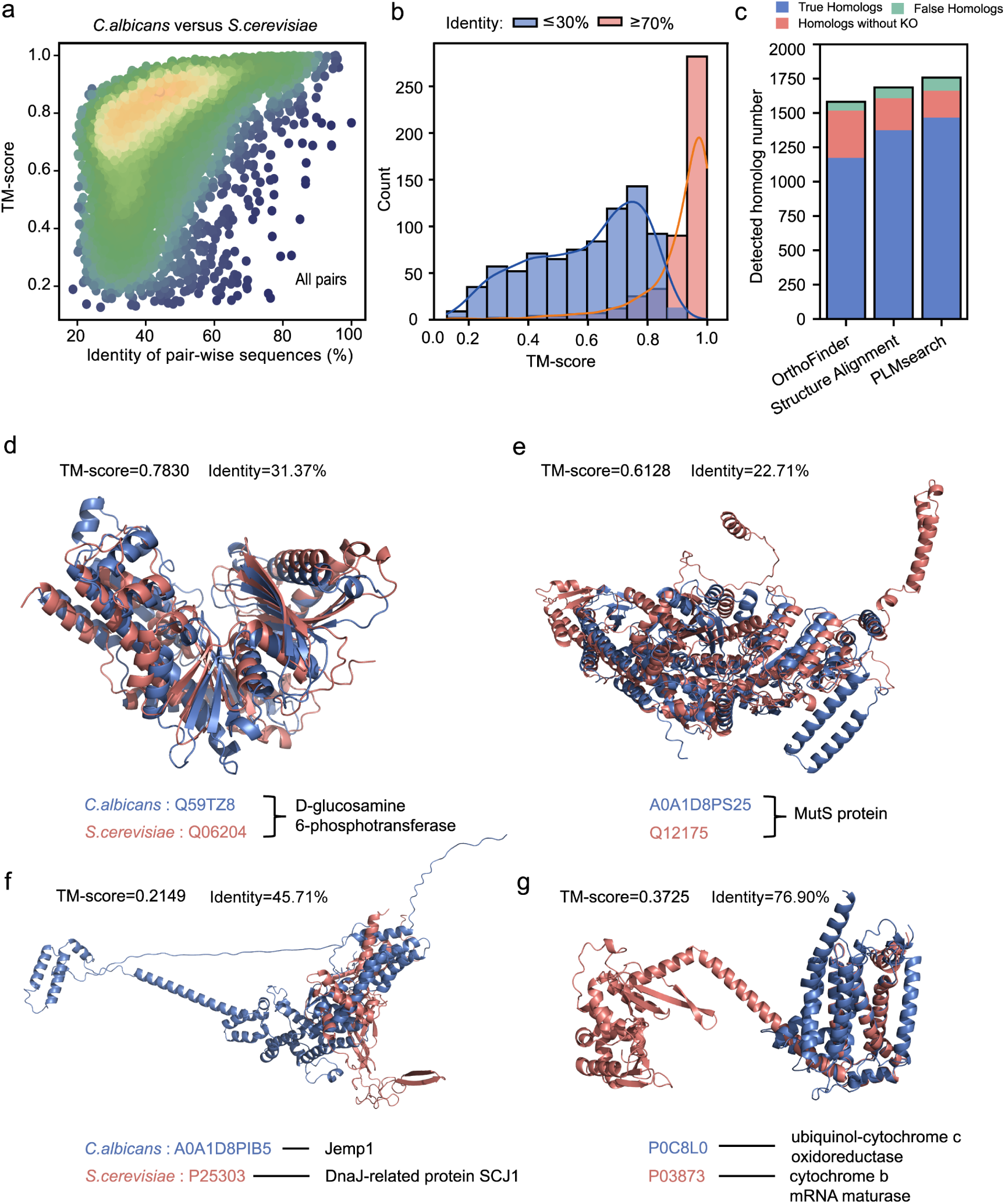
AlphaGEM enhances the inference of protein homologous relationships between two distinct species by combining sequence alignment and structure alignment. Distribution plots of sequence identity from sequence alignment and TM-scores from structural alignment between *C. albicans* and *S. cerevisiae* (a). Distribution of TM-scores for gene pairs with high sequence identity (>70%) and low sequence identity (<30%) (b). Accuracy in inference of protein homology using three procedures: the alignment based on sequencee blast, the alignment integrating both sequence and structural information; and the alignment based on PLMSearch, one protein language model. Here, the homologous relationships annotated by KEGG Orthology (KO) groups was used as the reference (c). Protein pairs with low sequence identity (OrthoFinder non-clustered) but high structural similarity (TM-score > 0.6) could still have the same catalytic functions (d, e). Protein pairs with high sequence identity but low structural similarity (TM-score < 0.4) were accompanied with divergent functions (f, g).

Notably, for more distantly related species and complex organisms such as mammals, increasing the threshold used in protein structure alignment based on Foldseek could improve accuracy in identifying protein homologous relationships, while still keeping a large number of homologous pairs between two organisms (Methods, Supplemental Fig. 2e), thus the structure-based protein function inference can be as a better approach than the sequence-based procedure in refining the protein homologous relationships between two chosen species.

### Protein language model accelerate the inference of homologous relationships between proteins across species without protein structure comparison

To ensure the broad applicability of AlphaGEM on the condition that the computational resource is limited, particularly for large-scale analyses involving hundreds of species, we developed a novel procedure for homologous relationships inference with the aid of PLMSearch^24^ (Methods, Fig. 1a, Supplemental Fig. 1b). This alternative strategy mainly uses protein language models to detect homologous protein pairs without relying on structural data. Taking *C. albicans* as an example (Fig. 2c, Supplemental Fig. 3a), this strategy achieved accuracy comparable to the above integrative approach accounting for both sequence and structure similarity (93.86% vs. 94.71%) while identifying additional homologous pairs (6.61%) missed by the latter.

To improve the reliability of the new strategy, we introduced a post-filtering step after the homologous relationships inference by PLMseacrh, in which the sequence coverage (>0.6 in *C. albicans* and >0.7 in mouse) and identity (>30% in *C. albicans* vs. >45% in mouse) acquired with blastp^30^ were used to refine the output of PLMSearch (Methods, Supplemental Fig. 1b) to balance sensitivity and specificity (Supplemental Fig. 3) in identifying homologs. For instance, excessively stringent identity thresholds (>50% in *C. albicans*) led to a sharp decline in valid homologous pairs, while overly lenient thresholds increased false positives. What’s more, we found that a few undetected homologous relationships by PLMseacrh can be recovered with the classical bidirectional best hits (BBH) identified through global BLAST alignment. Notably, species-specific parameter setting was critical, for example, the sequence divergence in mammalian genomes (e.g., *M. musculus*) required higher identity thresholds to consistent with structural alignment outcomes (Supplemental Fig. 4b). Taken together, alternative homologous relationships inference using PLMSearch could be resource-efficient for high-throughput GEMs reconstruction, whereas structure-based methods remain preferable for precision-focused tasks when computational resources permit.

### Improved annotation for nonhomologous proteins using an ensemble pipeline

Accurately annotating nonhomologous proteins in the input organism is an important step in identifying species-specific metabolic pathways in order to refine GEMs. Traditional approaches, such as eggNOG-mapper, are extensively used for annotating unknown proteins, but suffer from poor precision^26^. Relying on the breakthroughs in deep learning, several methods have shown much better prediction performance than traditional approaches^27^. However, using a single procedure for protein function prediction frequently generates unreliable results, which may lower the quality of the species-specific GEMs. To address this, AlphaGEM incorporates an ensemble pipeline integrating CLEAN, DeepECtransformer, eggNOG-mapper, and PLMSearch for comprehensive protein annotation to identify potential catalytic enzymes (Fig. 3a). To ensure robustness, we allocated weight scores to annotation results based on each tool’s reliability and specificity. Tools with high sensitivity but low accuracy were given a lower weight score, whereas those with higher specificity received a higher weight score. For instance, PLMSearch, while effective in identifying correct homologous relationships between two organisms, might introduce a significant number of false positives when used for protein function annotation, resulting in lower F1-scores (Fig. 3b, 3c). Thus, a lower weigh score was assigned to the output of PLMSearch when annotating the protein function with our ensemble pipeline. The sum of weight scores calculated from those four methods were used to rank the annotation results for each enzyme. We evaluated the annotation results using proteomics data from *C. albicans* and *S. pombe* that had been manually curated and partially experimentally validated. Accuracy analysis of protein function annotations revealed that the F1-scores of our ensemble pipeline (0.804, 0.410) were higher than those of individual tools (Fig. 3b, 3c): CLEAN (0.654, 0.326), DeepECtransformer (0.423, 0.189), PLMSearch (0.105, 0.097), and eggNOG-mapper (0.137, 0.094). It displays that our integrative pipeline could improve the enzyme function prediction in both F1-score and precision when compared to any single procedure, demonstrating the reliability of our pipeline. Subsequent evaluation shows that the annotation F1-score increased with the number of integrated tools (Fig. 3d, 3e). Note that the chosen best threshold in total weight score used in AlphaGEM could be adjusted to enhance the annotation performance of nonhomologous proteins according to the input organisms (Fig. 3f, 3g).

**Figure 3.**
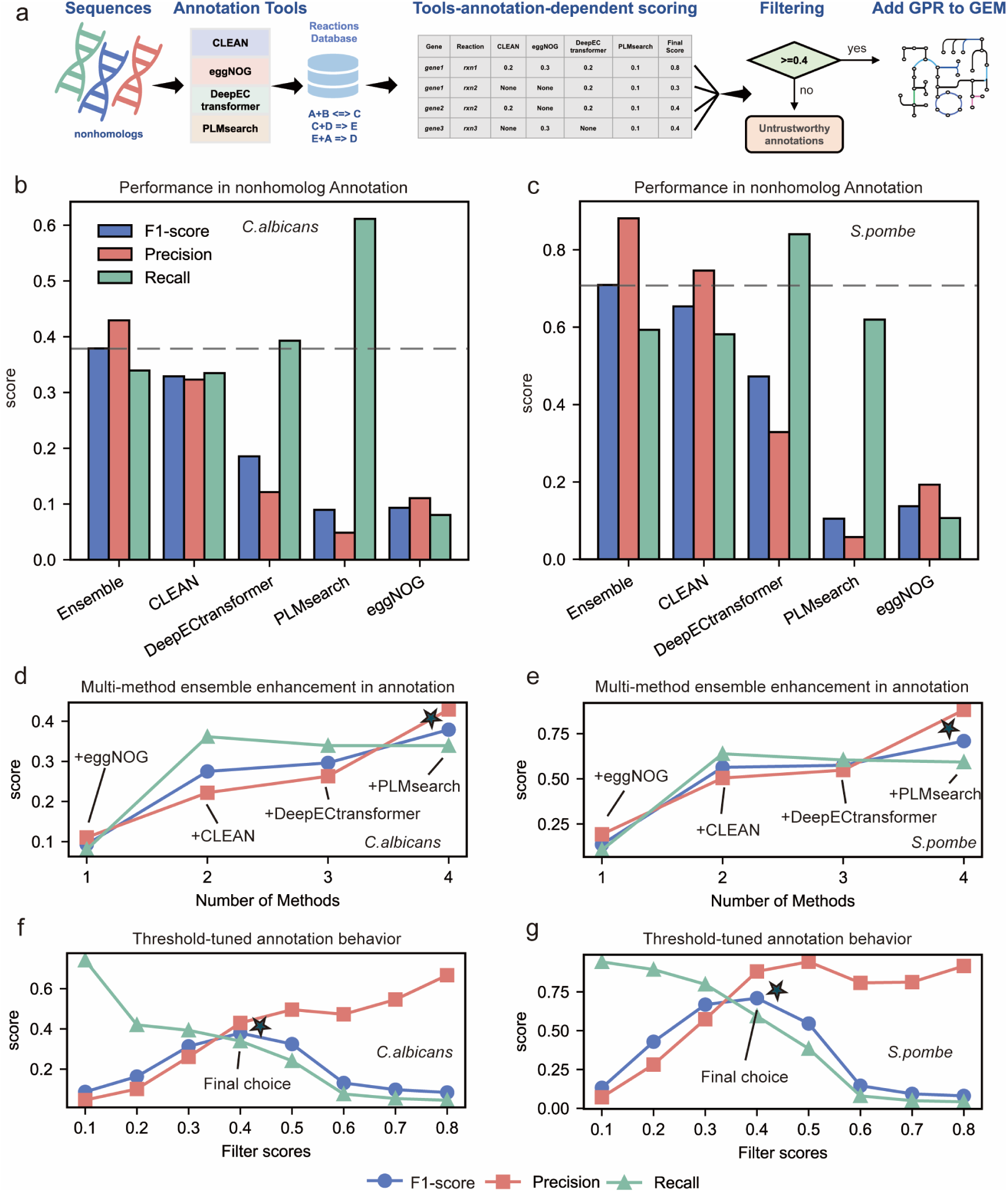
Systematic annotation of nonhomologous proteins in AlphaGEM. For nonhomologous proteins, AlphaGEM executes parallel functional annotation using CLEAN (EC numbers), DeepECtransformer (EC numbers), PLMSearch-based function search (Rhea reactions), and eggNOG-mapper (KO). All annotations are mapped to Rhea reactions, and each protein-reaction pair is assigned a cumulative confidence score based on tool-specific weights: CLEAN (0.2), DeepECtransformer (0.2), PLMSearch (0.1), and eggNOG-mapper (0.3). Pairs with aggregate scores ≥0.4 are retained; their gene-protein-reaction (GPR) rules are reconstructed and integrated into the draft GEM (a). Performance comparison between individual annotation tools and the ensemble method used in this work in nonhomologous protein annotation for *C. albicans* (b) and *S. pombe* (c). Profiles of annotation performance when incrementally adding tools into the ensemble method when conducting the nonhomologous protein annotation for *C. albicans* (d) and *S. pombe* (e). Influences of filtered score on the performances in nonhomologous protein annotation for *C. albicans* (f) and *S. pombe* (g).

### Comprehensive evaluation of AlphaGEM in generating ready-to-use GEMs

To systematically evaluate the performances of AlphaGEM in generating ready-to-use GEMs, we firstly constructed GEM for *C. albicans* and *S. pombe*, two widely used model yeast species.

We assessed the prediction capabilities of these models in prediction of carbon source utilization and essential genes, respectively.

In order to probe how each step within AlphaGEM framework influences model output, we built models for *C. albicans* and *S. pombe* using homologous pairs generated at different stages: 1) OrthoFinder-based orthology mapping, 2) US-align structural filtering, and 3) Foldseek-based structural expansion (Methods). Our analysis revealed that US-align structural filtering could reduce the coverage of proteins in the GEMs of *C. albicans* and *S. pombe*. The subsequent Foldseek-based structural expansion partially recovered the coverage of proteins. Crucially, while US-align filtering minimally affected the counts of metabolic reaction in two GEMs, the Foldseek based structural expansion enhanced metabolic reaction coverage. Finally, our ensemble pipeline enabled high-throughput protein function prediction, through which a few more genes, reactions and metabolites could be merged into GEMs (Supplemental Fig. 4a, 4b, 4c).

**Figure 4.**
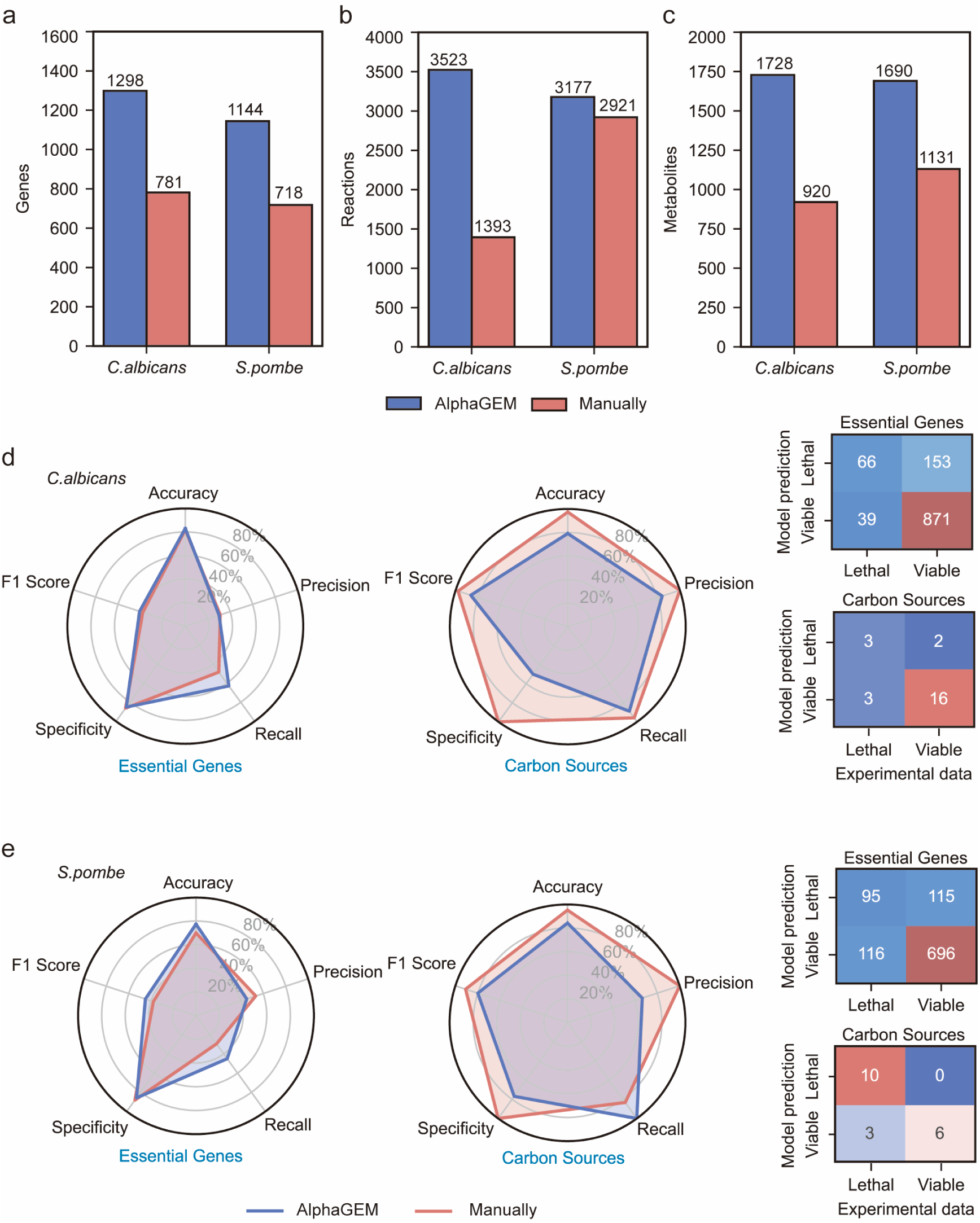
Quality assessment of GEMs for two simple eukaryotes reconstructed by AlphaGEM. Comparison in number of genes (a), reactions (b), metabolites (c) between GEMs of *C. albicans* (*S. pombe)* from manual curation and AlphaGEM. Comparison in accuracy for predicting carbon source utilization and essential gene by different GEMs from manual curation and AlphaGEM for *C. albicans* (d) and *S. pombe* (e).

Through the above steps, the final version of GEMs for two yeast species were generated and compared to the manually curated models^34, 35^. It shows that our models demonstrated significantly increased coverage in levels of genes, reactions, and metabolites (Fig. 4a, 4b, 4c). By further comparison, we found that *C. albicans* GEMs reconstructed by AlphaGEM performed well in predicting the usage of single carbon source (accuracy = 79.46%) and essential genes (accuracy = 82.99%) (Fig. 4d), yielding findings equivalent to a manually curated model^34^ (97.44% and 81.94%, respectively). Notably, AlphaGEM outperforms the manual reconstructed models in essential gene prediction. By contrast, our model performs somewhat worse than the manually curated model in predicting carbon source utilization, which likely due to the fact that the manually curated model has undergone iterative manual correction for carbon source utilization, thus enhancing the model prediction accordingly. Specially, we discovered that the GEMs built by AlphaGEM could predict the phenotypes of *C. albicans* using xylitol and D-glucosamine without introducing additional false-positive predictions. The enzymes implicated in the major pathways for these two carbon sources were structurally recognized as homologous proteins to *S. cerevisiae* enzymes (D-glucosamine 6- phosphotransferase and xylitol transporter), but no homologous proteins were discovered using sequence-based clustering (Fig. 2d). This demonstrates that our strategy could, to some extent, enhance the reliability of GEMs by refining the protein function annotation inferred solely by sequence-based analysis. Similarly, the GEMs built by AlphaGEM for *S. pombe* also achieved comparable accuracy in predicting carbon source utilization and essential genes (84.21%, 77.39%) compared to manually curated counterpart (95.24%, 69.89%) (Fig. 4e). While manually curated models benefit from iterative manual curation in carbon source predictions by comparing that with experimental dataset, AlphaGEM demonstrates comparable performance in generating high-quality GEMs for downstream applications.

### AlphaGEM enhance the reconstruction of high-quality GEMs for both bacteria and mammals

To further demonstrate AlphaGEM’s performance across diverse organisms, we constructed GEMs respectively for bacteria and mammals (Fig. 1a) respectively, and compared them with models generated by one of the state-of-art toolboxes and manually curated counterparts.

As gapseq is one of the best toolboxes for creating bacterial metabolic models and performs better than CarveMe and modelseed^19^, we comprehensively assess the quality of models generated by gapseq and AlphaGEM. *K. pneumoniae* is a ubiquitous human pathogen that can cause pneumonia, blood-spreading infections, urinary tract infections, and other life-threatening infections. We generated GEMs for *K. pneumoniae* with gapseq and AlphaGEM, respectively, and the models were compared to an experimentally validated model, iYL1228^36^. The efficacy of these three models for predicting carbon-source utilization was compared to experimental datasets. AlphaGEM achieved a prediction accuracy of 75.32%, comparable to the manually curated model iYL1228 (81.8%) (Fig. 5a). In contrast, the model constructed by gapseq exhibited poor specificity (16.67%), indicating that reactions and pathways added by gapseq were accompanied with excessive false positives, leading to poor substrate utilization specificity and functional deficiencies. In terms of model content, the AlphaGEM-constructed model annotated more genes (1,579) compared to gapseq (1,260) and iYL1228 (1,228) (Fig. 5c). AlphaGEM annotate more genes than other approaches, because the *K. pneumoniae* genome has 5728 genes, while the *E. coli* genome consists of 4401 genes, and the *E. coli* model iML1515 has 1515 annotated metabolic genes. The lower number of genes annotated by gapseq may be due to its less accurate gene functional annotation method, with the evidence that gapseq identified 63 reactions during gap filling while AlphaGEM only required 2 reactions to fill gaps, which suggests that the model constructed by gapseq suffers from insufficient gene annotation, resulting in more reactions required for gap filling. However, the AlphaGEM model contained fewer reactions (3,068) and metabolites (2,475) than the gapseq model (3,207 reactions and 2,577 metabolites), likely due to gapseq’s tendency to incorporate entire pathways from databases such as MetaCyc^37^, and the model it constructed exhibits an excessive number of genes corresponding to each reaction and an overabundance of reactions encoded by each gene, which results in the phenomenon of having fewer genes but more reactions (Supplemental Fig. 5a, 5b). This often introduces redundant reactions and metabolites, resulting in poor phenotype prediction, consistent with the phenomenon that gapseq-based GEM has a low specificity in predicting carbon source utilization (Fig. 5a). We further constructed GEMs for *B. subtilis* as the second case study and benchmarked it against models generated by gapseq and manual curation^38^. As GEMs for this species have not been extensively benchmarked for single carbon source utilization^38^, we validated model performance in essential gene predictions. AlphaGEM demonstrated prediction accuracy comparable to gapseq (AlphaGEM=81.87%, gapseq=80.30%), though it still weakly lagged behind the manually curated GEM (accuracy=89.26%) (Fig. 5b). Together, these results indicate that AlphaGEM performs comparably to manually curated models and, to some extent, outperforms the existing toolboxes in bacterial modeling.

**Figure 5.**
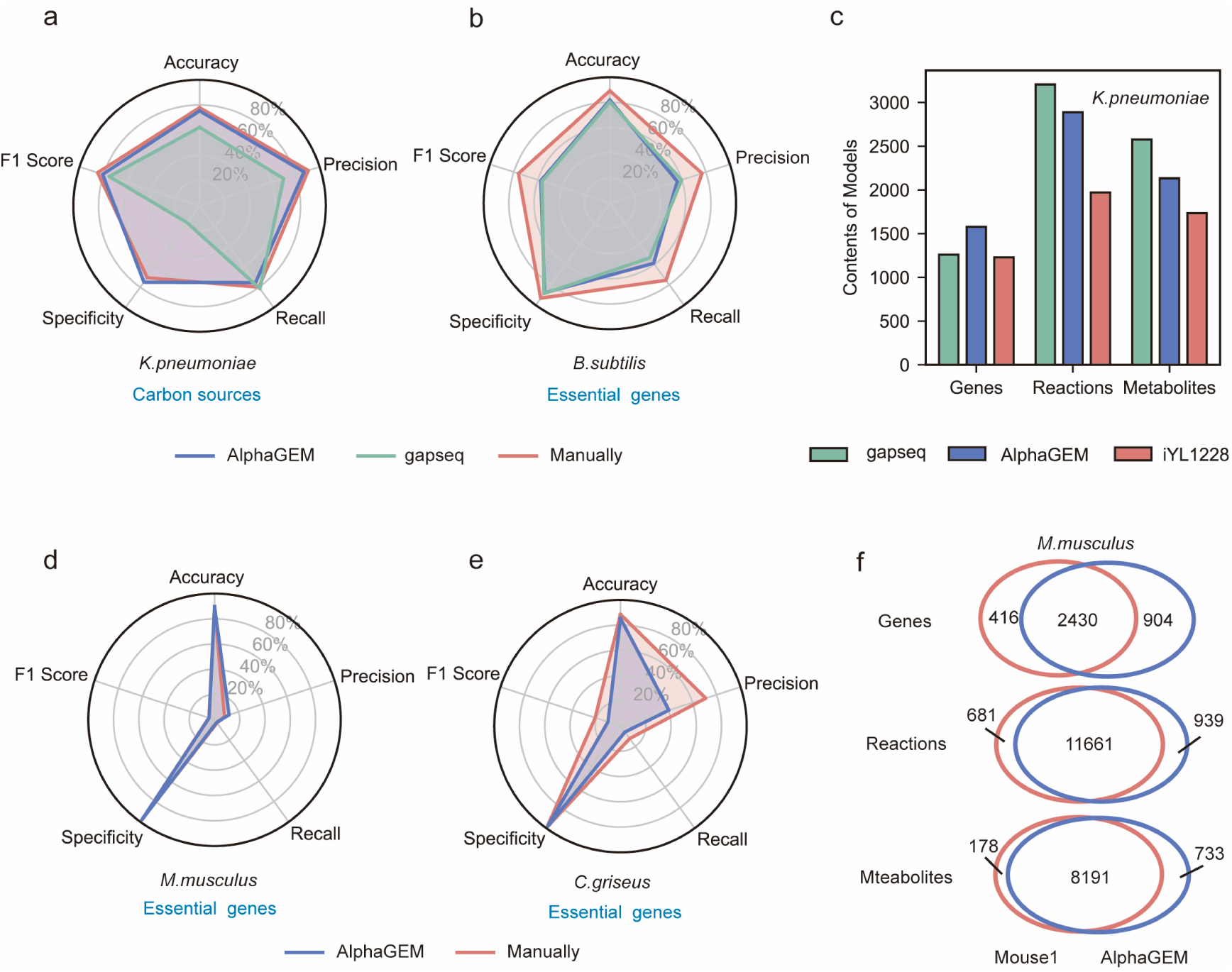
Quality assessment of GEMs for both bacteria and mammals reconstructed by AlphaGEM. Comparison in accuracy for predicting carbon source utilization by different GEMs from manual curation, gapseq and AlphaGEM for *K. pneumoniae* (a). Comparison in accuracy in essential gene prediction by different GEMs from manual curation, gapseq and AlphaGEM for *B. subtilis* (b). Comparisons in the number of genes, reactions, metabolites from different versions of GEMs for *K. pneumoniae* (c). Comparison in accuracy in essential gene prediction by different GEMs from manual curation and AlphaGEM for *M. musculus* (d) and *C. griseus* (e). Common and unique genes, reactions and metabolites analysis between two different GEMs from manual curation and AlphaGEM for *M. musculus* (f).

For mammals, after increasing the threshold for homologous relationship identification, we used the latest Human1 as the reference model to build GEMs for *M. musculus* and *Cricetulus griseus*, respectively, then compare the model’s performance with previously published ones. AlphaGEM achieved comparable essential gene prediction accuracy compared to the existing *M. musculus* GEM - Mouse1 ^39^ (model by AlphaGEM: F1-score = 4.32%, MCC = 1.84%, accuracy = 91.10%; Mouse1: F1-score = 4.71%, MCC = -0.16%, accuracy = 88.62%) (Fig. 5d). AlphaGEM’s multi-scale protein annotation approach, leveraging structural data for homology-based annotation, resulted in 488 additional annotated genes than Mouse1 (Fig. 5f) (Supplemental Fig. 2e) while the number of reactions and metabolites from the two types of models were comparable. To evaluate the accuracy of our annotations, we used multiple deep-learning-based annotation tools to mine the function of proteins corresponding to the unique genes from each model. We applied a higher threshold to filter these proteins, and those annotated with reactions after filtering were considered reliable metabolic proteins, while others were considered unreliable metabolic proteins. We found that, among the differential genes in Mouse1, 9.4% encoded reliable metabolic proteins, while 64.5% of the genes in the GEM constructed by AlphaGEM encoded considered reliable metabolic proteins (Supplemental Fig. 6a, 6b). Here, if we select only the genes encoded reliable proteins as the target GEM’s genes, the number of genes in the constructed model (3013) will be similar to that of Human1 (2887), which further supports the validity of our approach. For models constructed for *C. griseus*, AlphaGEM demonstrated accuracy and specificity in essential gene prediction comparable to manually curated model^40^ (accuracy 85.32% compared to 88.56%, specificity 98.52% compared to 99.29%). However, due to the limited number of essential genes identified through our approach, the F1-score performance was lower (10.43% compared to 21.09%) (Fig. 5e). For higher mammals, the complexity and abundance of intracellular pathways, coupled with relaxed flux constraints, can lead to essential reactions appearing as non-essential for growth, whereas the fact that some metabolic reactions are not being uncovered in the model would result in the opposite outcome. Adding additional constraints, such as transcriptomic data and enzymatic constraint, could improve prediction accuracy, further enhancing its usability for complex organisms^41^. Overall, these findings initially demonstrate that AlphaGEM could be leveraged to generate functional, comprehensive GEMs for mammals.

### AlphaGEM accelerates the generation of GEMs at large scale

The advancement of high-throughput sequencing technology has led to a large amount of genomic information from non-model organisms. Traditional procedures for generation of GEMs for these non-model organisms often undergo multiple rounds of iterative upgrades, require significant time and resources^42^. In this aspect, AlphaGEM could enable high-throughput reconstruction of GEMs to bridge the gaps between cellular phenotypes and genotypes at large scale. To confirm this, we leveraged AlphaGEM to automatically reconstruct GEMs for 332 sequenced yeast species, which were previously modeled using manually curated GEMs^43, 44^. By clustering the models using t-SNE based on their reaction content (Fig. 6a), we found that models from AlphaGEM exhibited distinct clustering patterns, in which the corresponding yeast species from the same genus displaying comparable aggregation with manually created ones, thus, to some extent, reflecting species classification based on phylogenetic analysis^44, 45^. Statistical validation using the Kruskal–Wallis test strongly supports this observation, confirming significant clustering structure in AlphaGEM (x: *P* value = 6.42 × 10^-27^, y: *P* value = 2.46 × 10^-35^, z: *P* value = 5.37 × 10^-46^) and manually created (x: *P* value = 4.33 × 10^-31^, y: *P* value = 1.09 × 10^-28^, z: *P* value = 2.68 × 10^-45^). Furthermore, models constructed by AlphaGEM exhibited greater hamming distances between reaction profiles compared to manually curated ones, indicating GEMs built by AlphaGEM could reflect species-specific metabolic differentiation (Fig. 6b).

**Figure 6.**
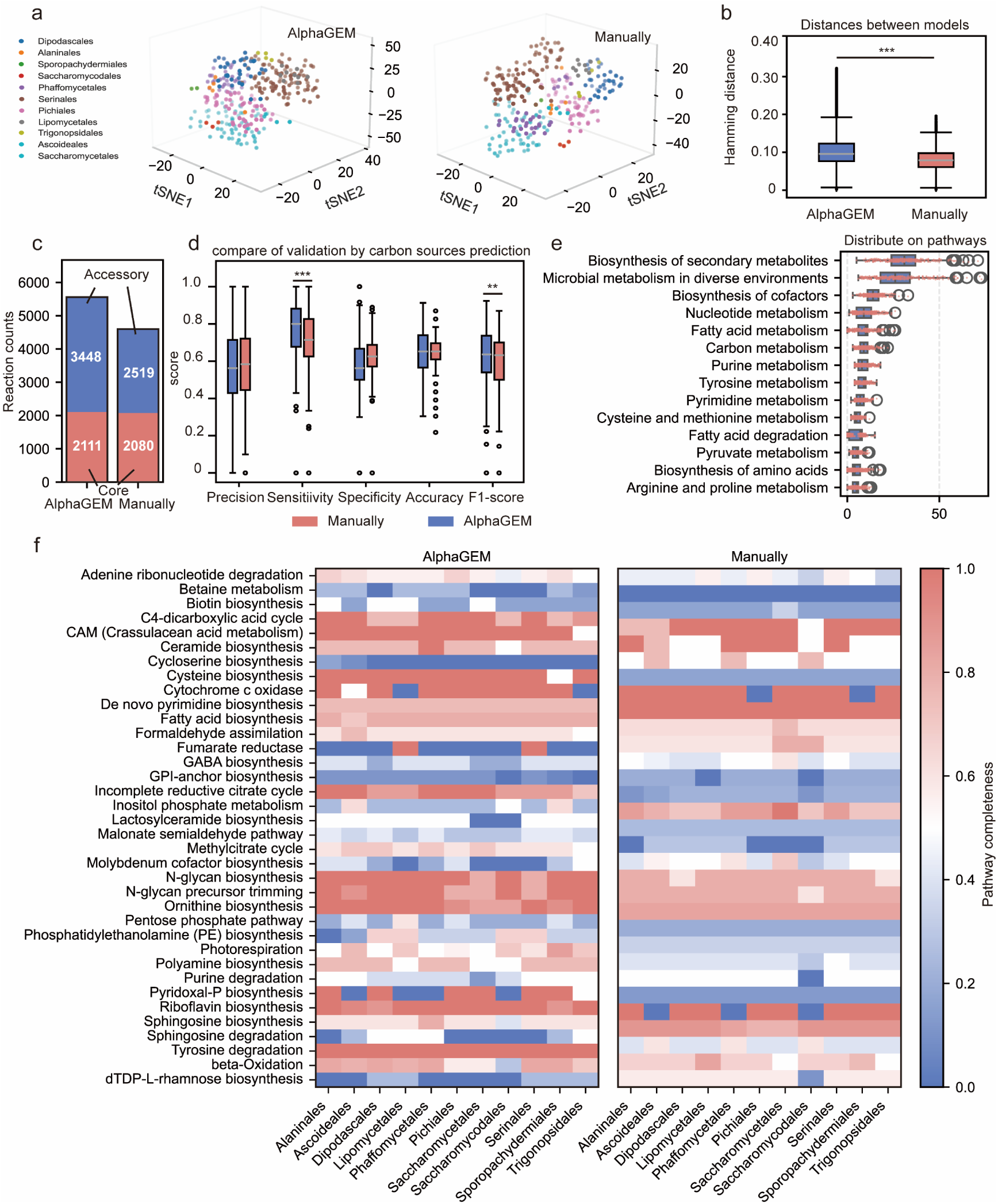
AlphaGEM accelerates high-throughput modeling for 332 distinct yeast species. The t-SNE clustering analysis of yeast species based the reaction existence in the manually curated GEMs and GEMs built by AlphaGEM, with different colors representing different genera (a). Comparison of Hamming distances between 332 yeast-species-specific GEMs constructed by AlphaGEMs and manually curated GEMs (b). Distribution of pan-, core-, and accessory-reactions across 332 yeast species (c). Validation in prediction of carbon source utilization using GEMs constructed by AlphaGEM and the manually curated GEMs (d). Main subpathways with newly added reaction only occurred in GEMs generated by AlphaGEM compared to manually curated GEMs (e). Heatmap of pathway completeness between GEMs constructed by AlphaGEM and the manually curated GEMs, focusing on which have top mean inter-genera differences. Color intensity indicates completeness values (red: 1.0; blue: 0) (f). (*: *P* value < 0.05; **: *P* value < 0.01; ***: *P* value < 0.001)

We compared structural characteristics between AlphaGEM-generated and manually curated models. There exist linear correlations (R² > 0.5) between two types of models in numbers of gene and metabolite (Supplemental Fig. 7a, 7b). While the number of reactions showed comparatively weak correlation (R² = 0.29), AlphaGEM could identify more reactions for most yeast species (Supplemental Fig. 7c, 7f). These findings indicate that the models reconstructed by AlphaGEM could be comparable to manually curated counterparts. Further analysis of GEMs across 332 yeast species demonstrated remarkably similar core reaction sets (AlphaGEM: 2111 vs. manual: 2080) (Fig. 6c), confirming AlphaGEM’s accuracy in identifying conserved yeast metabolic functions. Notably, AlphaGEM annotated significantly more accessory reactions (3448 vs. 2519) (Fig. 6c), consistent with its enhanced species-specific metabolic differentiation observed earlier. This expanded accessory repertoire showcases AlphaGEM’s potential to uncover broader metabolic pathway diversity across distinct species.

To validate the functional quality of the models, we performed carbon source utilization predictions using the newly reconstructed GEMs and compared them with experimentally verified phenotypes (Fig. 6d). It shows that the GEMs built by AlphaGEM achieved comparable F1-score and accuracy compared with manually constructed ones. Additionally, the GEMs built by AlphaGEM were accompanied with a slight high sensitivity (*P* value < 0.01), indicating comparable performances in predicting specific carbon source utilization by GEMs from our toolbox and the reported ones. However, the specificity of GEMs built by AlphaGEM was slightly lower, which may be due to that the employment of PLMsearch could result in more false-positive homologous relationships when generating GEMs at large scale. In contrast, manually constructed GEMs, which were corrected based on experimental phenotypes, exhibited higher accuracy for certain predictions.

Lastly, the AlphaGEM’s capability in uncovering dark metabolism was explored. Here, these reactions uniquely annotated by AlphaGEM in each species were mapped onto metabolic pathways. We found that the novel reactions identified by AlphaGEM were primarily distributed in pathways related to secondary metabolism, microbial metabolism in diverse environments, and cofactor biosynthesis (Fig. 6e). This distribution pattern demonstrates AlphaGEM’s potential to elucidate untapped metabolic capabilities within secondary metabolic pathways. Furthermore, we evaluated the potential of AlphaGEM for uncovering gaps in pathway metabolism by analyzing pathway completeness. We then evaluated pathway completeness to pinpoint gaps in metabolic networks, concentrating on those pathways showing the largest inter-genus differences between GEMs built by AlphaGEM and manual curated ones (Fig. 6f). In these pathways, AlphaGEM exhibited greater variability across genera, demonstrating its capacity to uncover dark metabolism in a genus-specific manner by tailoring reaction sets to each lineage rather than applying a one-size-fits-all approach.

## Discussions

In this work, we present AlphaGEM, an integrated toolbox for automatic, high-throughput construction of genome-scale computational models for target organisms, uniquely addressing the critical gap in modeling numerous newly sequenced species with high quality^46, 47^. Our tool fundamentally differs from existing methods (e.g., gapseq^19^) by achieving two breakthrough capabilities: (1) high-throughput generation of eukaryotic GEMs with quality approaching manually curated benchmarks and (2) extension to prokaryotic modeling with performance comparable or superior to state-of-the-art tools. This dual advantage stems from our integration of protein structural alignment and deep learning-based protein function inference, enabling precise detection of distant homologies widely present in understudied organisms. Importantly, we developed an ensemble algorithm that could expand metabolic network coverage for target organisms through the in-depth annotation of non-homologous proteins using multiple latest deep learning models. To demonstrate these capabilities, we systematically assess the performance of AlphaGEM in reconstruction of GEMs across diverse domains, including fungi (332 different yeast species), prokaryotes (e.g., *K. pneumoniae*), and mammalian cells (e.g., *M. musculus*), demonstrating AlphaGEM’s excellent capability for large-scale, multi-species metabolic characterization and revealing its potential for uncovering dark metabolism in less characterized organisms through enhanced annotation of non-canonical enzymatic functions.

Although our strategy offers a robust framework for developing high-quality GEMs for non-model organisms, several challenges still remain. Firstly, generating all protein structures for one species requires substantial computational resources, which may lower the efficiency of our pipeline as a whole. Additionally, using protein language models, i.e., PLMSearch, could identify homologous relationships at large-scale, but, to some extent, compromise accuracy at the same time. Moreover, the predictive accuracy of deep learning models employed in our study could directly influence AlphaGEM’s effectiveness, as training datasets used in different deep learning models often contain inconsistencies, leading to potential errors in assigning key reactions to specific proteins. Future advancements in development of advanced computational tools and high-quality datasets will critical to address these limitations, thus further helping to enhance the overall performances of AlphaGEM.

In conclusion, AlphaGEM enables automatic, high-quality metabolic modeling across diverse species. It paves the way for systematic exploration of dark metabolism and support scalable, next-generation analysis of newly sequenced organisms, thereby advancing our understanding of metabolic processes in microbes and higher organisms on Earth.

## Methods

### Inference of Homologous relationships

In this study, two methods were employed to infer protein homologous relationships between target and reference species as the initial step in reconstructing draft GEMs for target organisms.

1) Homologs Based on Structural Alignment

Initially, OrthoFinder was utilized to cluster proteins from annotated genome assemblies, generating pairwise protein similarity matrices. Then, protein structure alignment was conducted for each protein pair using US-align, yielding a structural similarity score (TM-score). The results were filtered based on the TM-score and the predicted protein structure quality score (pLDDT)^48^. Protein pairs with a TM-score exceeding the threshold (default: 0.7) were classified as exhibiting high structural similarity, while those with a pLDDT below the threshold (default: 0.8) were excluded from structure alignment due to low-quality structure predictions (Supplemental Fig. 2d). These filtered protein pairs constituted the first proteins homology set (Proteins Homology Set One).

Foldseek was employed to conduct a structure-based homology search, expanding the collection of homologous protein pairs. Homologous pairs identified through Foldseek-based alignment were screened using the following criteria: a TM-score ≥ 0.8, coverage ≥ 0.9, and probability = 1 (Supplemental Fig. 2a, 2b). Pairs with a pLDDT below the default threshold of 0.8 were excluded due to low-quality structure predictions. Metabolism proteins were categorized into transporters and non-transporters, with transporters further filtered using a identity threshold of 30%. To enhance the accuracy of structure-based homologs identification, homologous pairs were clustered separately for each enzyme type. For clustering homologous proteins from the reference species, the DBSCAN algorithm (default eps = 1) was applied. Clusters were selected based on the highest combined TM-score and coverage values, ensuring all pairs meet the established TM-score and coverage thresholds. This process generated the second proteins homology set, referred to as Proteins Homology Set Two. Finally, the two homology sets were combined to determine the final protein homologous relationships.

2) Homologs Based on PLMSearch

The embeddings of the target strain and the reference strain genomes generated by a protein language model were integrated as input to predict homologous relationships using PLMSearch. Based on a predefined SS-predictor threshold (default = 0.8), the results were filtered by blastp with coverage >= 0.6 and identity >= 30%. Subsequently, bidirectional best hits (BBH) identified through global BLAST alignment performed by diamond^49^ was added to the final set of homologs (with identity >= 60%, coverage >= 0.5).

### Homologs filtering using ProtTrans

Our algorithm identified several proteins with an unusually high number of corresponding homologous proteins (i.e., > 6) from the input organism (Supplemental Fig. 8b, 8c), which is biologically implausible. To address these issues, we utilized ProtTrans, a protein language models^50^, to encode the pre-identified homologous proteins. We then applied Principal Component Analysis (PCA) for dimensionality reduction to refine these homologous relationships (Methods, Supplemental Fig. 8a). This approach effectively reduced the number of homologs for proteins with multiple relationships to fewer than three, minimizing false positives (Supplemental Fig. 8b, 8c). This strategy to prune the number of homologous proteins could be an optional choice to build GEMs with our toolbox. This approach proved critical for constructing the mouse GEM, successfully differentiating functions between Cytochrome P450 26B1 and Cytochrome P450 26C1 (Supplemental Fig. 8e), thereby enhancing essential protein prediction accuracy. The detailed method is as follow:

For each protein (RP) executing a specific reaction in the reference model and its homologous proteins (P1, P2, …), they were grouped together. If the group contains only one protein apart from RP, this step was skipped. For the prediction groups, ProtTrans was used to generate embeddings for the sequences of all proteins in the group. PCA was then applied to reduce the dimensionality of these embeddings to analyze the distances between different sequences. The distance between the reduced PCA coordinates was defined as:

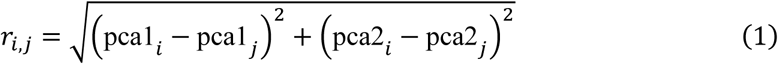

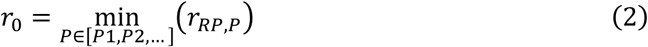

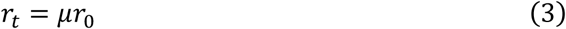

Where *i* and *j* represent different proteins within the group. The minimum distance *r*_0_ the reference protein RP was selected, and a threshold *r*_t_ was applied (default *μ* = 1.1). Homologous proteins with a distance greater than this threshold were removed, and the remaining group was considered the target homologous relationship.

### Validation of homologous protein relationship

KO numbers were assigned to proteins based on homologous protein relationships using data from UniProt^51^ and KEGG^52^. Proteins sharing the same KO number were designated true homologs; those with different KO numbers were designated false homologs. Accuracy was calculated as the proportion of true homologs among all evaluated protein pairs (true homologs / (true homologs + false homologs)).

### Selection of high-quality reference models

The latest published manually curated functional GEMs were selected as reference GEMs. For modeling eukaryotic single-cell organisms, the metabolic model of *Saccharomyces cerevisiae*, Yeast9^14^, is chosen as the reference model. Species with a closer phylogenetic relationship to *Saccharomyces cerevisiae* can use this model as a reference. For prokaryotic cells, the *Escherichia coli* model, iML1515^15^, is selected as the reference model, and species with a closer phylogenetic relationship to *E. coli* can also use it as a reference. The latest published human GEM, Human1^16^, is selected as the reference model for mammalian modeling. These reference models have been updated with metabolite and reaction IDs from the Rhea^53^ and ChEBI databases, using mapping relationships from MetanetX^54^ to ensure compatibility when adding new reactions. AlphaGEM further provides an interface for manual submission of custom reference models, which must be processed using the supplied curation script model_curation.py to align with framework requirements.

### Draft-Model reconstruction for non-model organisms

The GEMs for target species were reconstructed based on the homologous relationships derived from the reference models. First, reactions lacking homologous proteins in the reference model were removed according to the gene-protein-reaction (GPR) relationships. Subsequently, the retained reactions were re-annotated through homologs mapping to generate draft models. The proteins were ready to map the genes encoded themselves and were added to the GEMs.

For isoenzymes-catalyzed reactions, the reaction is considered to exist if at least one corresponding isoenzyme is present. The reaction GPR is defined based on the isoenzymes present. For reactions catalyzed by protein complexes, the reaction is considered to be present if over 80% of the complex’s proteins is identified, and the GPR for the reaction is reconstructed based on the complex. For transporters, an optional stricter filtering threshold is offered for their identification.

### Protein annotation through multiple computational tools

Proteins excluded from the homologous protein pairs were annotated using ensemble method. eggNOG-mapper^28^ was employed to annotate proteins with KO numbers, CLEAN^26^ and DeepECtransformer^27^ for EC number annotations, and Rhea IDs corresponding to UniProt protein data. CLEAN annotations were filtered at a confidence threshold of 0.7 before downstream analysis. PLMSearch^24^ identified homologous protein pairs and annotated target proteins with Rhea IDs by performing homology searches against UniProt’s Rhea-annotated proteins. The results are filtered using a threshold determined by SS-predictor (default = 0.9) to obtain the final output. All annotation results were standardized to RHEA IDs by rhea.rdf for consistent protein function annotation. The annotations are assigned with different weight score from different tools (CLEAN: 0.2, eggNOG-mapper: 0.3, DeepECtransformer: 0.2, PLMSearch: 0.1) (Fig. 4c). Annotations with weight ≥ 0.4 were selected as the final annotations.

### Validation of protein annotation through multiple computational tools

Standardized protein-to-reaction pairs were derived from UniProt Rhea^53^ annotations, serving as the benchmark for evaluating protein–reaction pairs annotated by computational tools. Accuracy was quantified using: 1) true positives (TP): protein-reaction pairs annotated by the tool that were also present in the database annotations; 2) false positives (FP): protein-reaction pairs annotated by the tool that were not present in the database annotations; 3) false negatives (FN): protein-reaction pairs present in the database annotations that were not annotated by the tool. These metrics (TP, FP, FN) were then used to calculate precision, sensitivity and F1-score. Notably, FP assessments exhibit inherent limitations due to incomplete reaction annotations in UniProt^51^.

### GEMs extension from rhea database

To accelerate reaction integration, biochemical reactions from Rhea^53^, a standardized reaction-metabolite database were retrieved. These reactions were converted into a pool of ready-to-use cobra.Reaction objects. All reactions and metabolite information were extracted from the rhea.rdf file, including the mapping relationships with other databases, reaction reversibility, metabolite composition, charge, and CheBI IDs^55^, to generate reaction classes that can be used with cobrapy. Reactions were subsequently added to the model based on prior Rhea ID annotations for genes.

### Improving the quality of GEMs using gap-filling

As the last step, the constructed model underwent gap-filling using the gap-filling function cobra.flux_analysis.gapfill() in cobrapy, with the reference model serving as the universal model. Gap-filling was optionally performed using either minimal or complete growth media. At the same time, the objective reactions (e.g., biomass production) and the biomass composition was consistent with reference GEMs. To further refine and filter potential gap reactions identified through gapfilling, AlphaGEM performed tBLASTn searches against the genome of the target organism using the proteins catalyzing these gap reactions. Homology-based replacement was then applied: for hits with a bit score > 200, the protein corresponding to the hit with the highest bit score was used to replace the original homologous protein in the reaction. If no qualifying hit (bit score > 200) was found, AlphaGEM attempted to remove the reaction from the model, while ensuring the integrity of the objective reaction. As a result, gap-filling further complemented GEMs by filling essential pathway gaps (e.g., only 3 reactions for *C. albicans*)

### Essential genes prediction

To conduct gene essentiality analysis, the essential gene list from published researches^34, 35, 38–40^ was used, in silico growth with gene essentiality was calculated as follows: 1) Each gene in the model was iteratively knocked out using the model.genes.knock_out() function. 2) Growth rates were simulated using FBA under identical minimal glucose medium conditions. 3) A gene was classified as computationally essential if the simulated growth rate was less than 1×10⁻⁴ h⁻¹.

Predictions were compared against the experimental essentiality datasets using a confusion matrix, categorizing results as: 1) true positive (TP): experimental non-essential / in silico non-essential; 2) true negative (TN): experimental essential / in silico essential; 3) false negative (FN): experimental non-essential / in silico essential; 4) false positive (FP): experimental essential / in silico non-essential (essential genes with in silico lower growth was calculated as negative).

### Carbon source utilization prediction

To conduct carbon source utilization analysis, the phenotype data from published researches ^34–36, 39^ was used, in silico growth with carbon source utilization was calculated as follows: 1) the model medium was set to the minimal growth medium with glucose replaced by alternative carbon sources. 2) Growth rates were simulated using FBA. 3) A carbon source was classified as no-growth if the simulated growth rate was less than 1×10⁻^3^ h⁻¹.

Predictions were compared against experimental observations using the same confusion matrix framework (TP, TN, FN, FP) as described for essential gene prediction (growth as positive and no-growth as negative).

### Metrics to assess the prediction performance of GEMs

To evaluate model performance, the following metrics were computed based on the confusion matrix, consisting of true positives (TP), true negatives (TN), false positives (FP), and false negatives (FN).

Matthews Correlation Coefficient (MCC)^56^:

The MCC, a balanced measure of classification quality, was computed as:

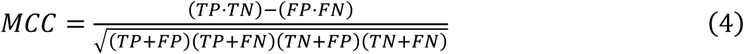

Sensitivity (Recall)^57^:

Sensitivity, the ability to correctly identify positive samples, was computed as:

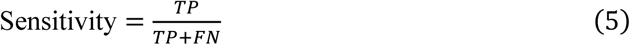

Accuracy^57^:

Accuracy, the proportion of correctly classified samples, was calculated as:

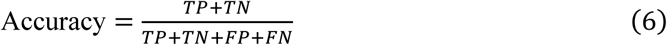

Specificity^57^:

Specificity, the ability to correctly identify negative samples, was computed as:

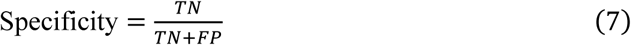

Precision^57^:

Precision, the proportion of true positives among predicted positives, was calculated as:

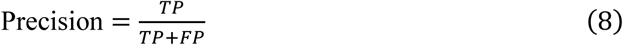

F1-Score^57^:

The F1-score, the harmonic mean of precision and sensitivity, was calculated as:

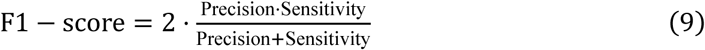

### Construction of GEMs using gapseq

Starting from the genome of *K. pneumoniae* and *B. subtilis* in UniProt, gapseq’s four-step default parameters were utilized to build the GEM, with the exception that during gap-filling, the medium was set to minimal medium to ensure the strain could grow under minimal conditions.

### Construction of GEMs for 332 Yeast Strains

We constructed Genome-Scale Metabolic Models (GEMs) for the annotated yeast genomes using AlphaGEM’s PLMSearch method under default parameters. Gap-filling was performed using a minimal medium formulation. Carbon source utilization predictions followed the method described in the preceding section. The resulting models were analyzed directly without modification. For dimensionality reduction, t-SNE (t-Distributed Stochastic Neighbor Embedding) was applied using the SciPy library^58^. This cluster analysis was conducted based on reaction presence/absence profiles.

### Comparison of Inter-genus pathway completeness differences

Reactions from AlphaGEM-constructed GEMs and manually constructed GEMs were extracted and mapped to KEGG-defined pathways. The pathway completeness for a single species is calculated as:

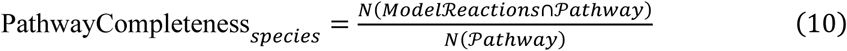

The genus-level assessment is defined as:

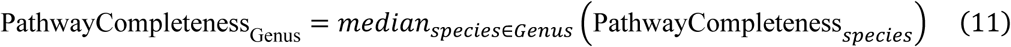

Where *N* denotes the cardinality of the reaction set, and median is computed over all valid species annotations in the target genus.

## Statistical analysis

1) T-test *P* value calculation

A two-tailed independent t-test was performed to determine the statistical significance of differences between two groups. For each pair of datasets ** the *P* value calculated using the formula:

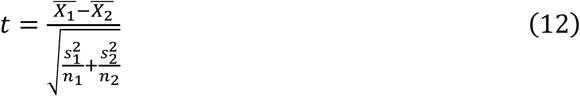

where 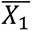 and 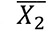 are the means of the two groups, 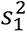 and 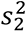 are the variances, and *n*_1_ and *n*_2_ are the sample sizes. The resulting t-statistic was used to compute the *P* value, which determines statistical significance at a predefined threshold (e.g., *P* value < 0.05).

2) Kruskal-Wallis test

The Kruskal-Wallis was performed test on the target t-SNE dimension (x or y or z coordinates) across all genus. The Kruskal-Wallis test statistic H is calculated as:

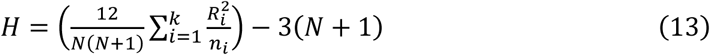

where N is the total number of samples, k is the number of genera, *n_i_* is the number of species in the i-th genus, and R_i_ is the sum of ranks of the i-th genus. Statistical significance was determined using a threshold of *P* value < 0.0001.

The upper statistical analysis calculations were performed using Python (v3.12) with the SciPy and Scikit-learn libraries.

## Code and data availability

The yeast genome data used in this study were sourced from ^44^, and manually curated GEMs for 332 yeast species were obtained from ^43^ .The source code for AlphaGEM is available on GitHub at https://github.com/hongzhonglu/AlphaGEMs.

## Declaration on the use of artificial intelligence tools for language refinement

During the preparation of this work, the authors utilized ChatGPT (OpenAI), DeepSeek-R1 (DeepSeek), and Grok-3 (xAI) exclusively for language editing and grammatical correction. All AI-processed text was critically reviewed, fact-checked, and revised by the authors. The authors assume full responsibility for the validity, integrity, and originality of the published content.

## Author Contributions

H.L conceived the study. W.H, L.X, and H.W. contributed to the development of AlpahGEM. All authors read, edited, and approved the final manuscript.

## Acknowledgments

This work is supported by grant 2022YFA0913000 from the National Key R&D Program of China, Shanghai Municipal Science and Technology Major Project, and grant 22208211 and 22378263 from the National Natural Science Foundation of China (NSFC). The funding body has no role in the design of the study, analysis and interpretation of the data, preparation of the manuscript, and decision to submit the manuscript for publication.

---

**Supplemental Figure 1.**
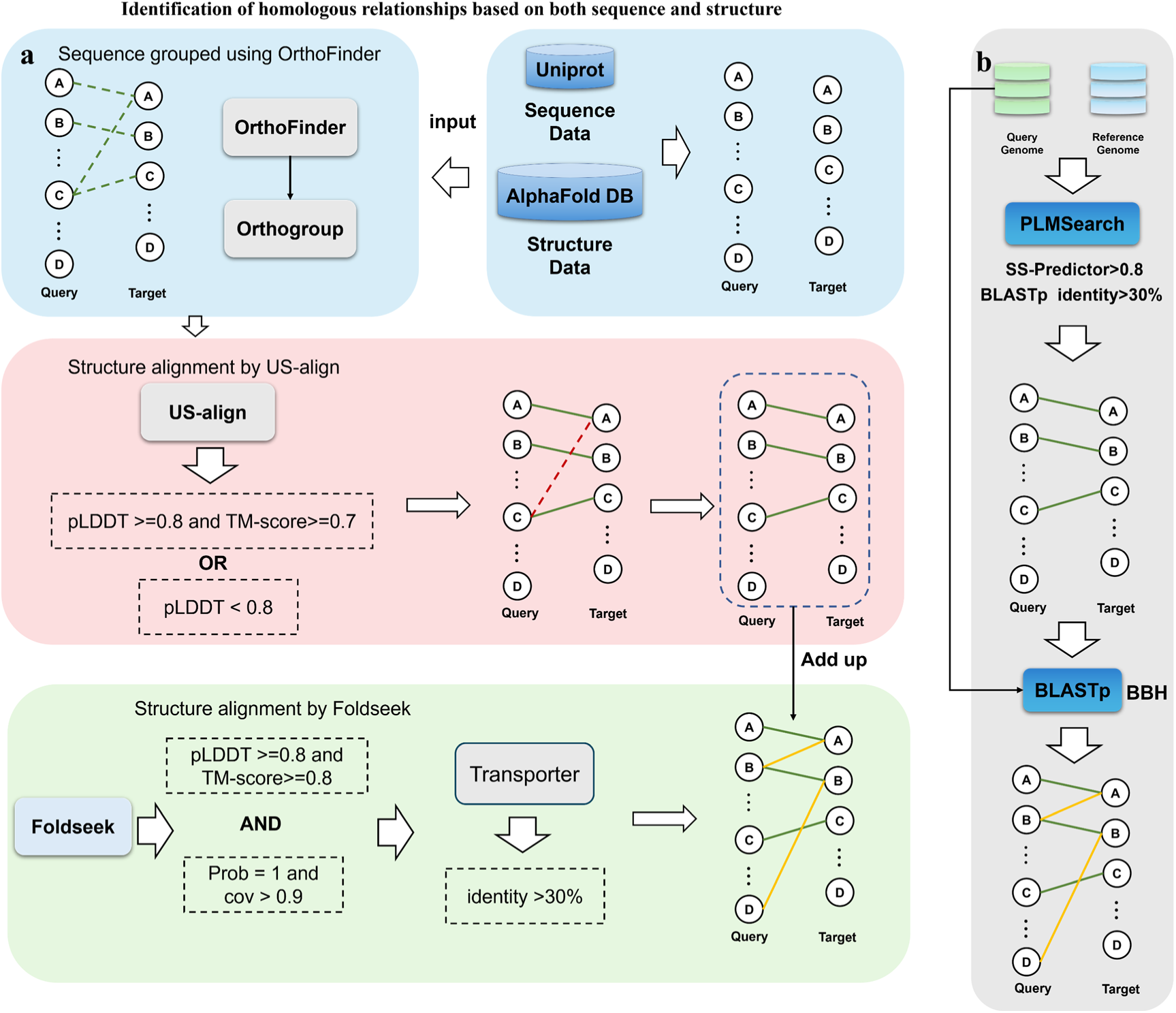
Homologous relationships identification process. Initial homologous relationships detection via sequence-based clustering (OrthoFinder), followed by structure-based filtering with USalign (TM-score/pLDDT thresholds) to eliminate false homologs, and global structural alignment-based homologous relationships expansion using Foldseek (a). Comprehensive homologous relationships search through PLMSearch with SS-predictor-parameter filtering, subsequent blastp verification of candidate pairs, and final BBH extension via global BLAST (b).

**Supplemental Figure 2.**
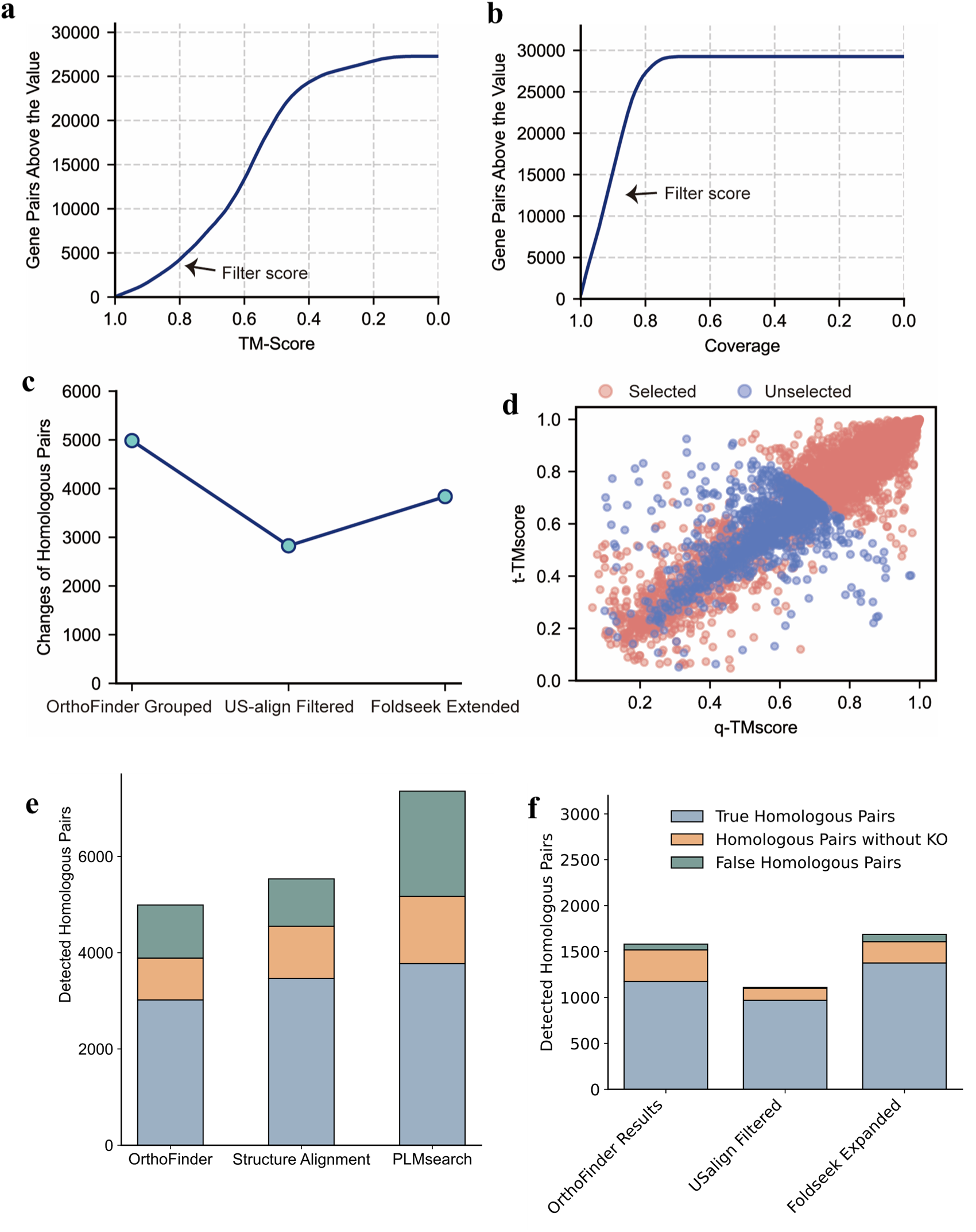
Changes in the number of homologous relationships during the modeling process. TM-score distribution and threshold selection for protein pairs between *C. albicans* and *S. cerevisiae* obtained from globally structural alignment using Foldseek (a), Coverage distribution and threshold selection for the same alignment (b). Changes of the number of homologous pairs across three modeling steps: OrthoFinder grouping, US-align filtering, Foldseek extension (c). q-TM-score and t-TM-score (TM-score normalized by the query/target length) distributions of structural alignment performed by USalign for sequence-based homologous pairs between *C. albicans* and *S. cerevisiae* (pLDDT and TM-score threshold selection) (d). Accuracy in inference of protein homology between *M. musculus* and *H. sapiens* using three procedures: the alignment based on sequence blast, the alignment integrating both sequence and structural information; and the alignment based on PLMSearch, one protein language model (e). Accuracy in inference of protein homology between *C. albicans* and *S. cerevisiae* by sequence clustering; integrated US-align-based structural filtering; integrated Foldseek-based structural extension (f).

**Supplemental Figure 3.**
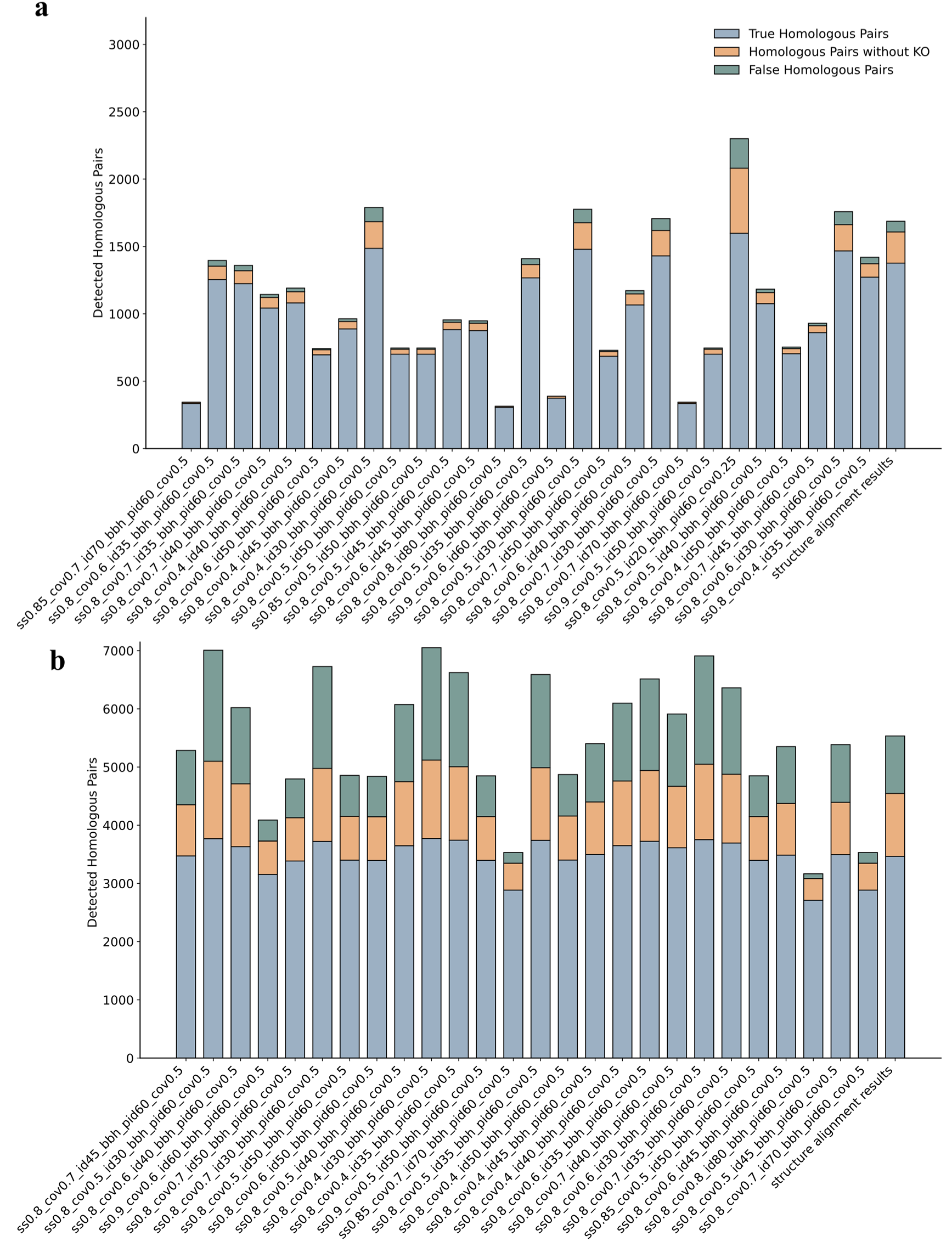
Comparison in protein homologous relationships detection between PLMSearch and structure-based annotation. Accuracy in inference of homologous protein pairs between *C. albicans* and *S. cerevisiae* identified by PLMSearch (with various blastp filtering threshold: coverage, identity) versus structural alignment (a). Accuracy in inference of homologous protein pairs between *M. musculus* and *H. sapiens* identified by PLMSearch (with various blastp filtering threshold: coverage, identity) versus structural alignment (b).

**Supplemental Figure 4.**
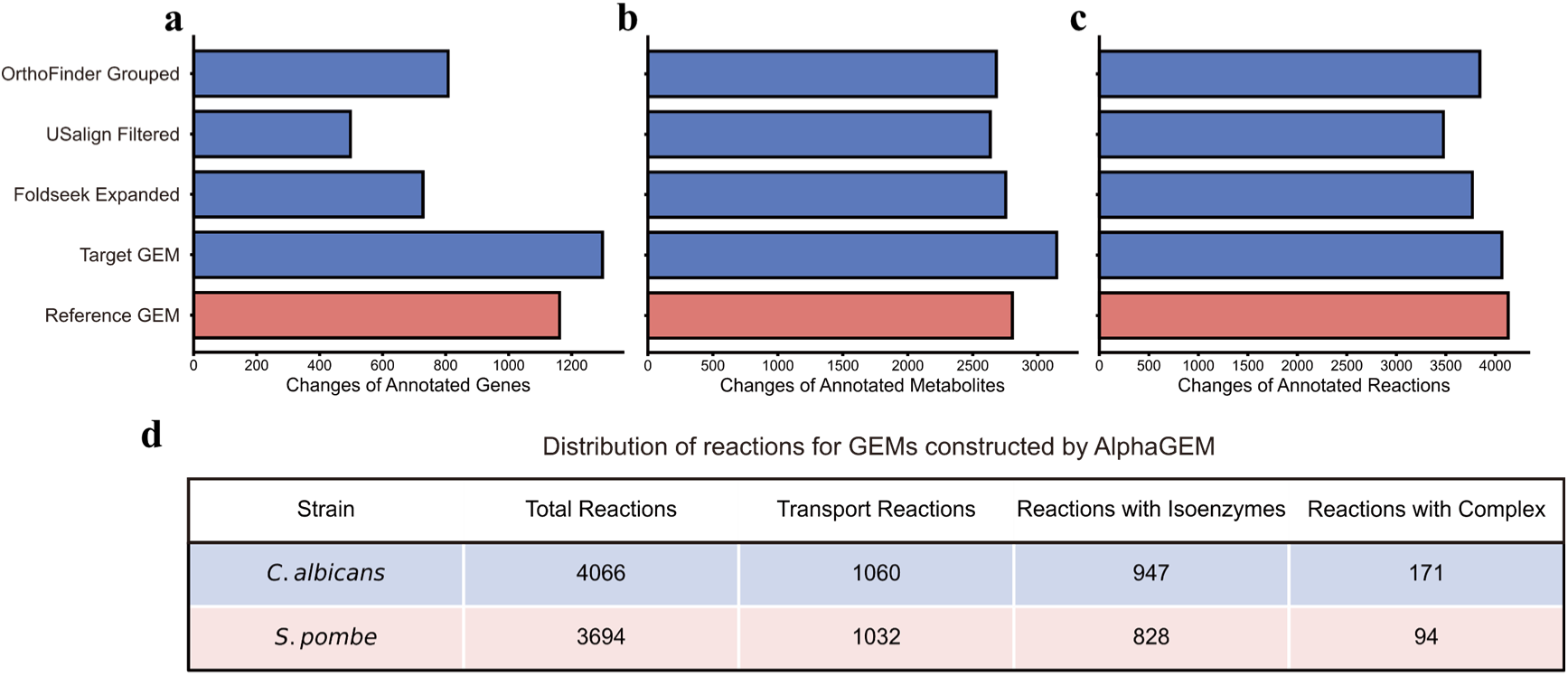
GEM reconstruction for *C. albicans* using AlphaGEM. Changes in the number of annotated genes (a), metabolites (b), and reactions (c) during the stepwise construction of the *C. albicans* GEM: OrthoFinder grouped; US-align Filtered; Foldseek expanded; final output GEM, along with the comparison with reference GEM: Yeast9. Distribution of different type of reactions in the constructed *C. albicans* and *S. pombe* models (d).

**Supplemental Figure 5.**
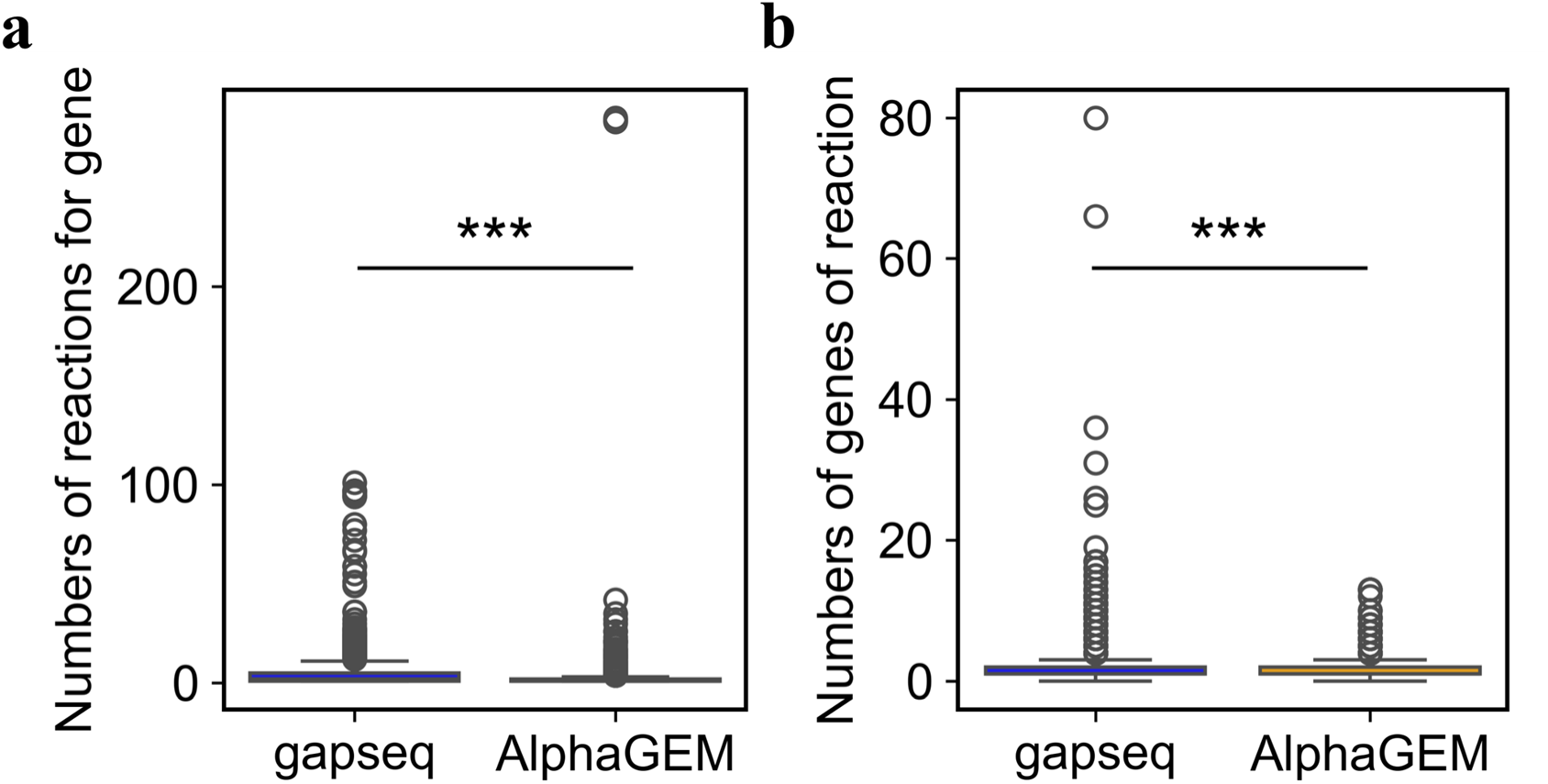
Comparison of *K. pneumoniae* GEMs constructed by gapseq and AlphaGEM, respectively. Distribution in the number of reactions catalyzed by one gene in GEMs built by gapseq and AlphaGEM, respectively, GEM built by gapseq has more extreme values (a). Distribution in the number of genes (including isozymes and enzyme complexes) catalyzing one reactions in GEMs built by gapseq and AlphaGEM, respectively, GEM built by gapseq has more extreme values (b). (***: *P* value < 0.001)

**Supplemental Figure 6.**
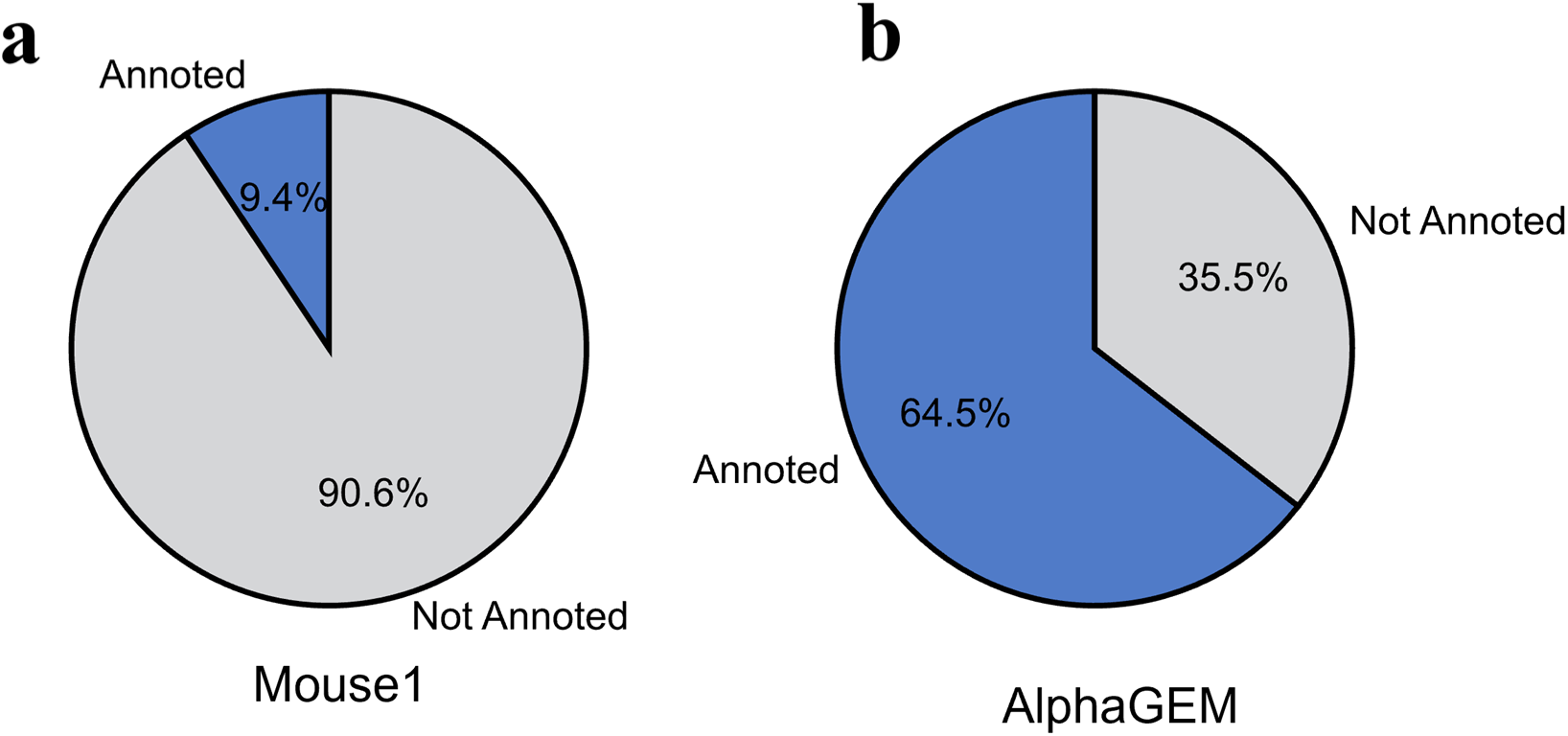
Comparison of differential gene-protein-reactions in Mouse1 and GEM reconstructed by AlphaGEM. Proportion of proteins annotated for different reactions in the Mouse1 model compared to AlphaGEM(a), in the AlphaGEM model (b). *Analysis performed by integrating multiple AI tools with a higher threshold (0.5) to identify differential reactions between models*

**Supplemental Figure 7.**
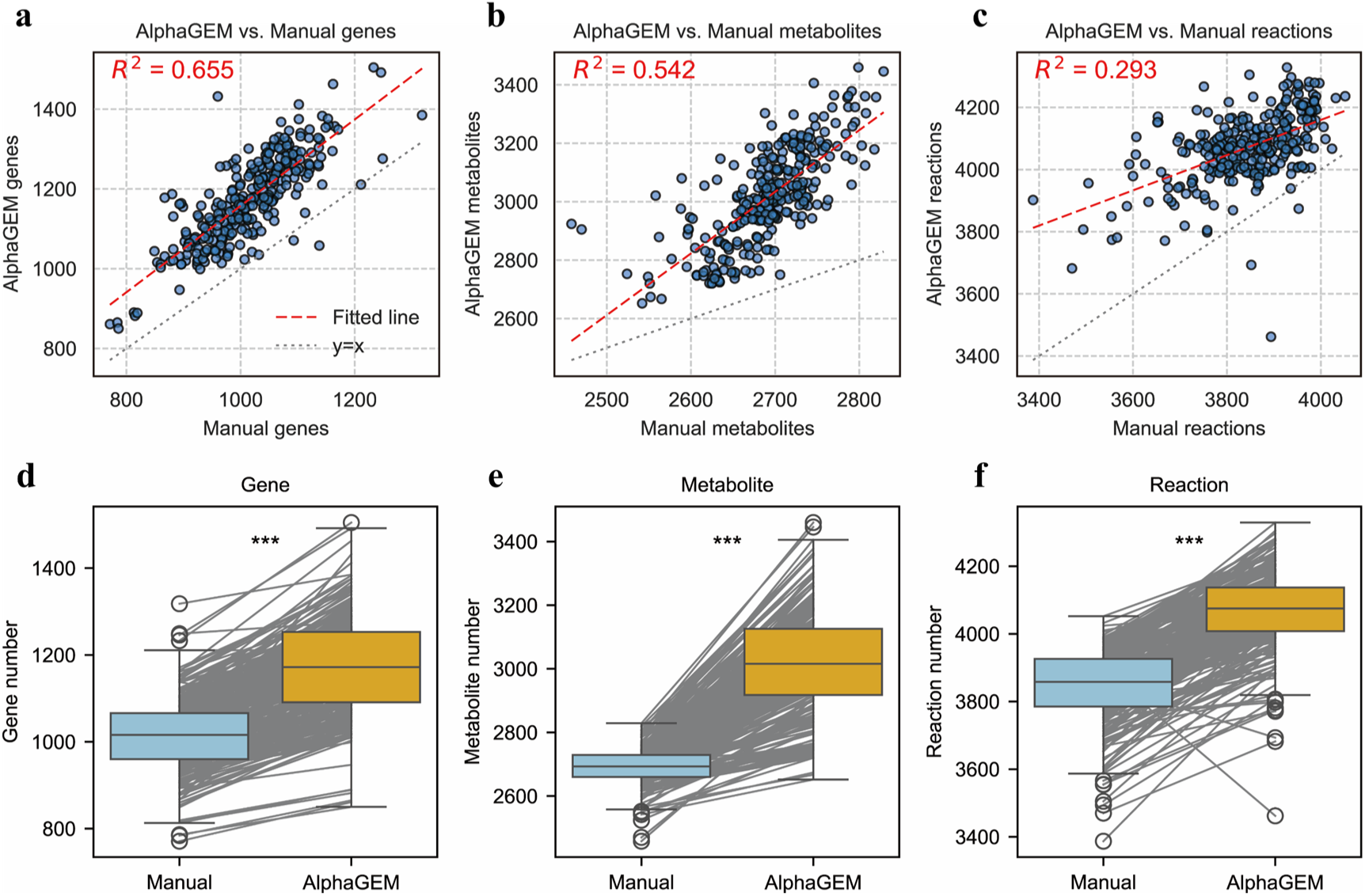
Comparison and linear regression analysis between GEMs built by AlphaGEM and manual curation for 332 yeast species. Correlation analysis in gene counts (a), metabolite counts (b), reaction counts (c) between GEMs built by AlphaGEM and manual curation. Comparison of gene counts (d), metabolite counts (e), reaction counts (f) between GEMs built by AlphaGEM and manual curation. (***: *P* value < 0.001)

**Supplemental Figure 8.**
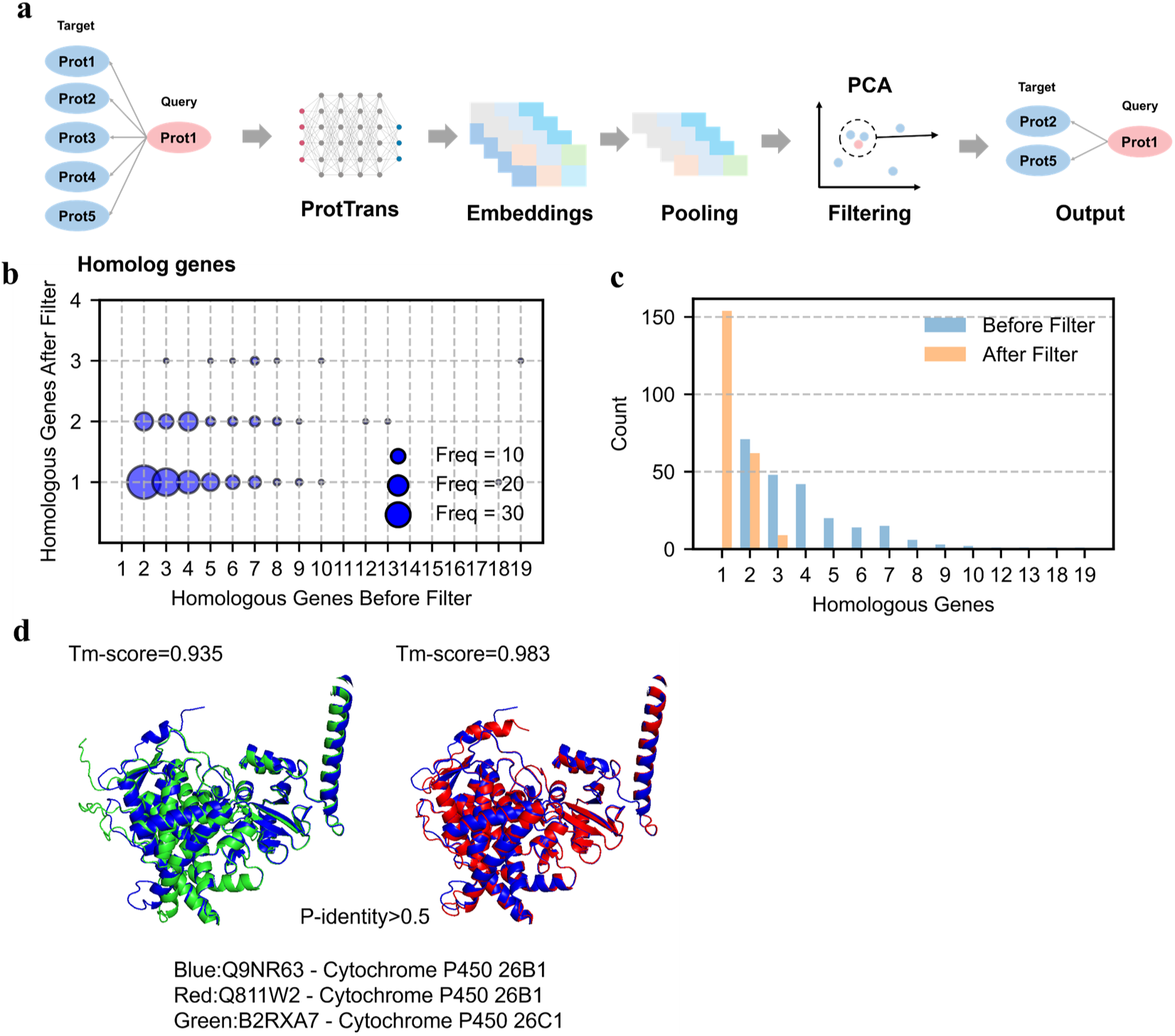
Homologous proteins could be refined using ProtTrans. ProtTrans-generated embeddings were used for proteins with multiple homologous proteins, followed by filtering based on a pre-selected distance threshold in PCA-reduced clustering space (a). Distribution of different changes in the number of homologous proteins per protein after filtering (b). Compare of global distribution in the number of homologous proteins per protein before and after filtering (c). This approach successfully eliminated erroneous homologous protein assignments derived from structural alignment during GEMs reconstruction (d).

