## Supplemental Figures for "AlphaGEM Enables Precise Genome-Scale Metabolic Modelling by Integrating Protein Structure Alignment with deep-learning-based Dark Metabolism Mining"

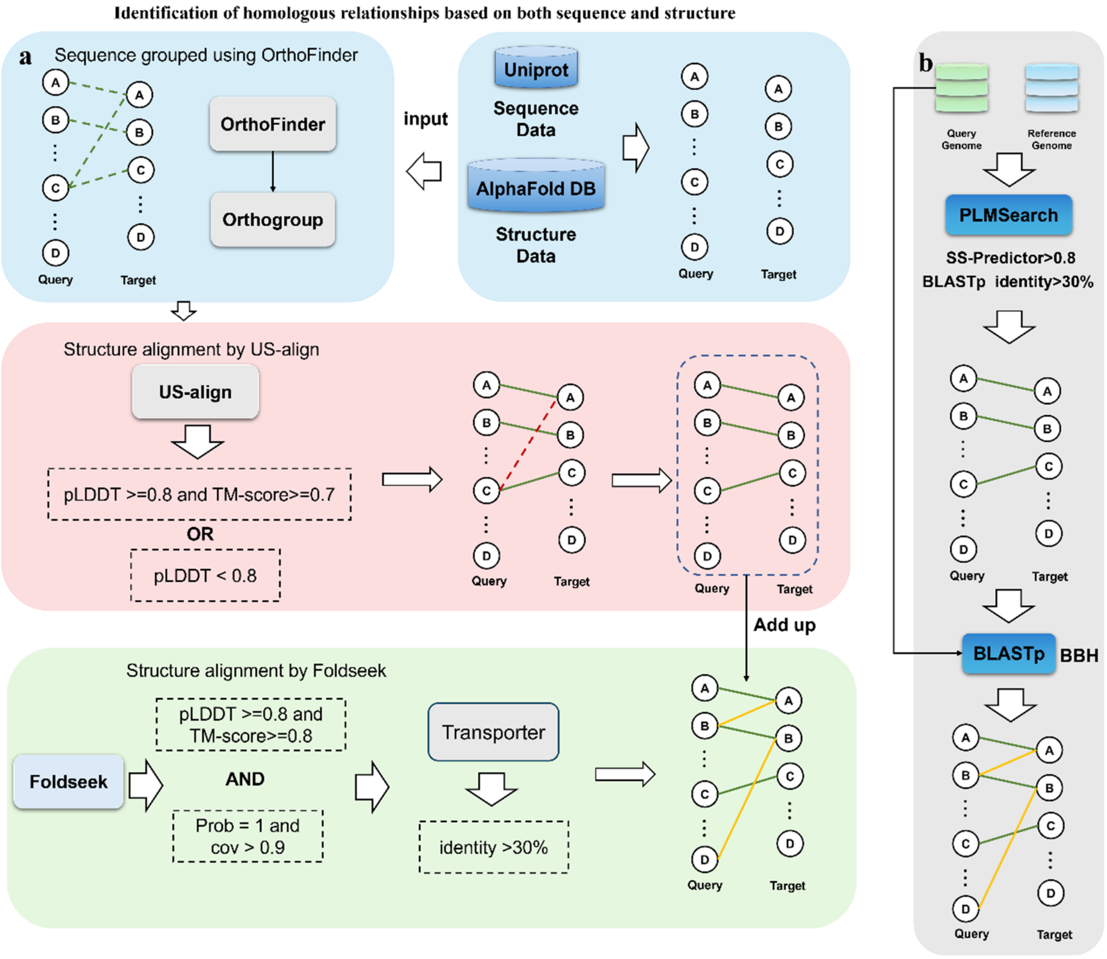


Supplemental Figure 1 Homologous relationships identification process.

Initial homologous relationships detection via sequence-based clustering (OrthoFinder), followed by structure-based filtering with USalign (TM-score/pLDDT thresholds) to eliminate false homologs, and global structural alignment-based homologous relationships expansion using Foldseek (a). Comprehensive homologous relationships search through PLMSearch with SS-predictor-parameter filtering, subsequent blastp verification of candidate pairs, and final BBH extension via global BLAST (b).

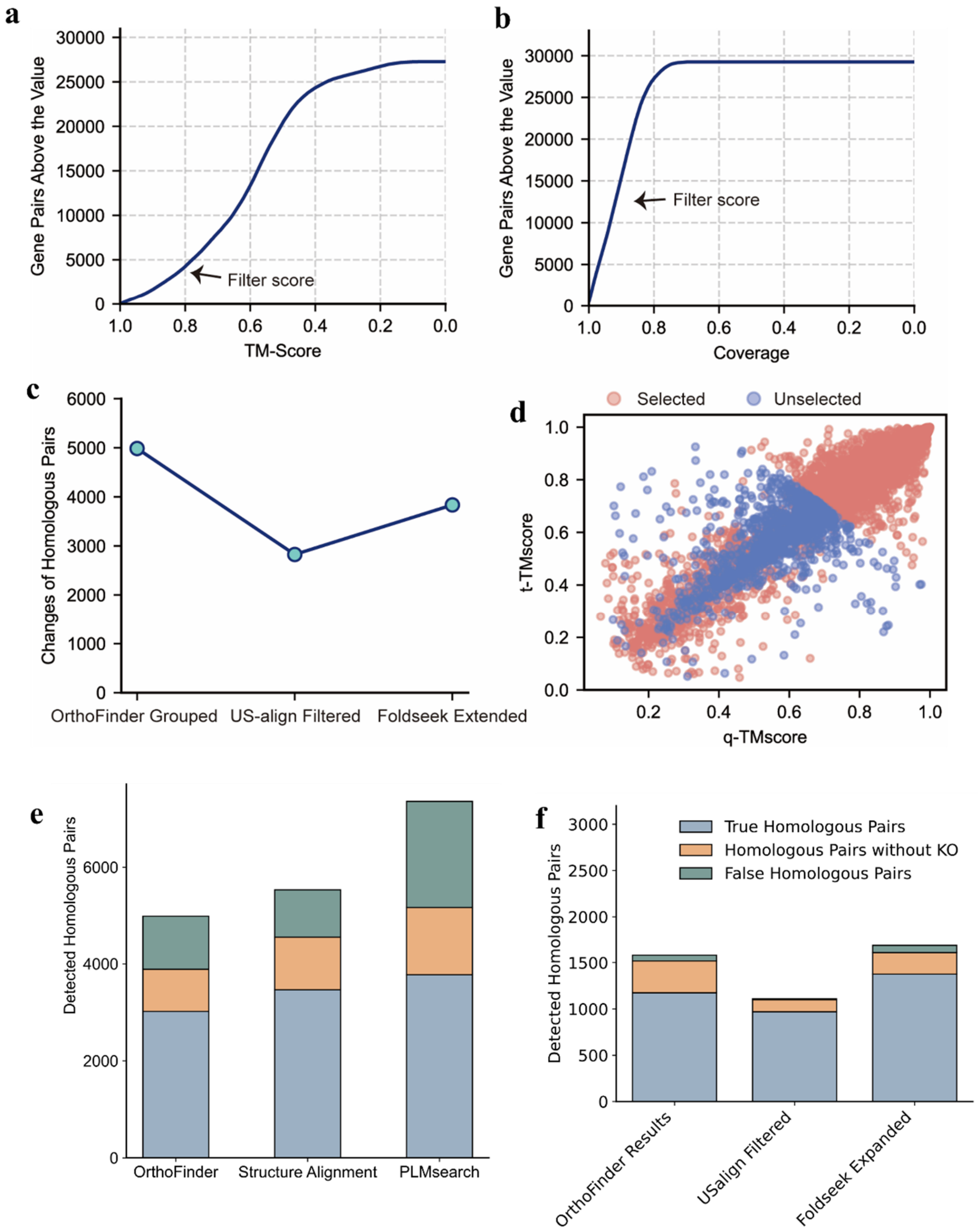


Supplemental Figure 2 Changes in the number of homologous relationships during the modeling process

TM-score distribution and threshold selection for protein pairs between *C. albicans* and *S. cerevisiae* obtained from globally structural alignment using Foldseek (a), Coverage distribution and threshold selection for the same alignment (b). Changes of the number of homologous pairs across three modeling steps: OrthoFinder grouping, US-align filtering, Foldseek extension (c). q-TM-score and t-TM-score (TM-score normalized by the query/target length) distributions of structural alignment performed by USalign for sequence-based homologous pairs between *C. albicans* and *S. cerevisiae* (pLDDT and TM-score threshold selection) (d). Accuracy in inference of protein homology between *M. musculus* and *H. sapiens* using three procedures: the alignment based on sequence blast, the alignment integrating both sequence and structural information; and the alignment based on PLMSearch, one protein language model (e). Accuracy in inference of protein homology between *C. albicans* and *S. cerevisiae* by sequence clustering; integrated US-align-based structural filtering; integrated Foldseek-based structural extension (f).

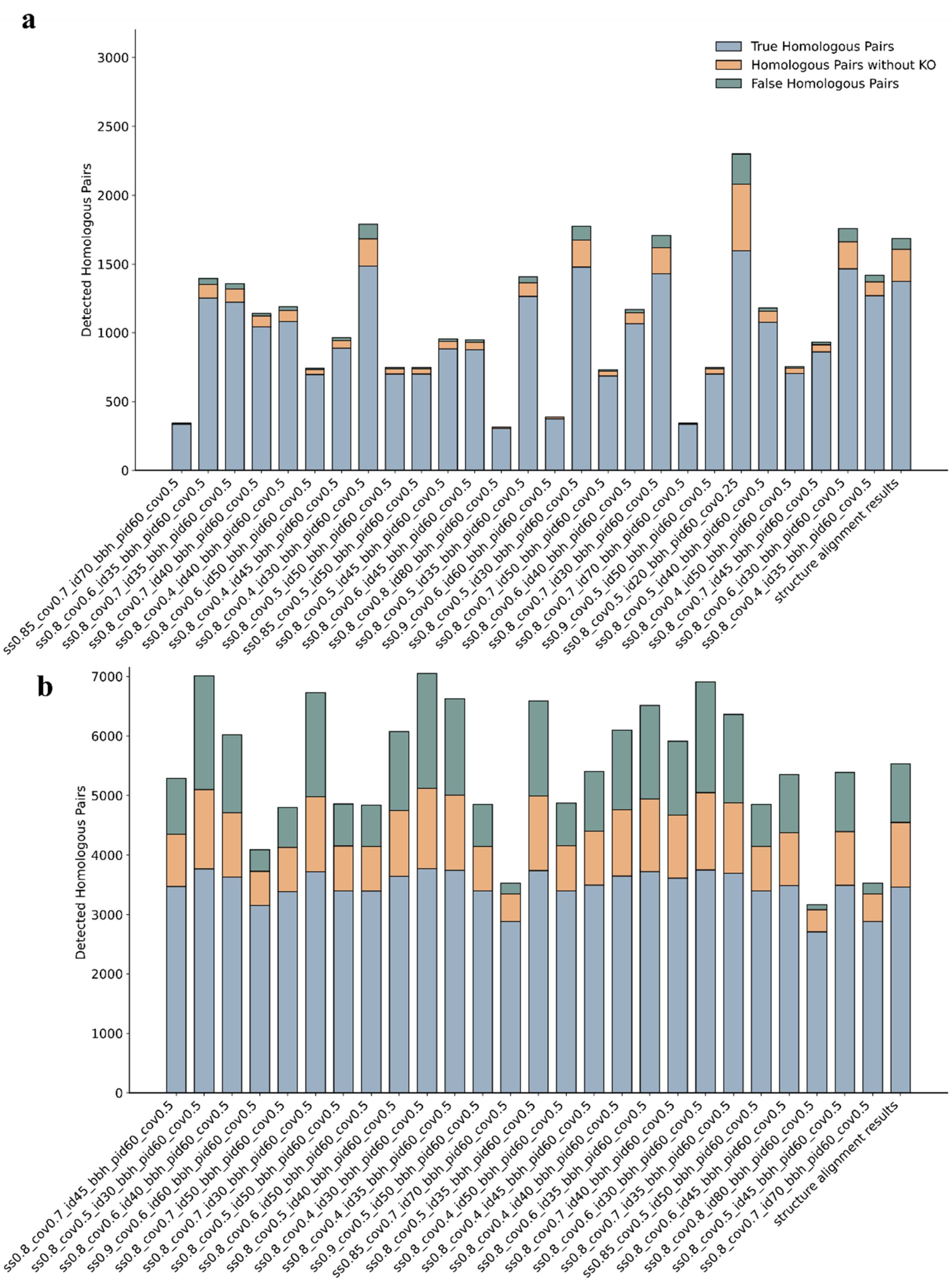


Supplemental Figure 3 Comparison in protein homologous relationships detection between PLMSearch and structure-based annotation.

Accuracy in inference of homologous protein pairs between *C. albicans* and *S. cerevisiae* identified by PLMSearch (with various blastp filtering threshold: coverage, identity) versus structural alignment (a). Accuracy in inference of homologous protein pairs between *M.*

*musculus* and *H. sapiens* identified by PLMSearch (with various blastp filtering threshold: coverage, identity) versus structural alignment (b).

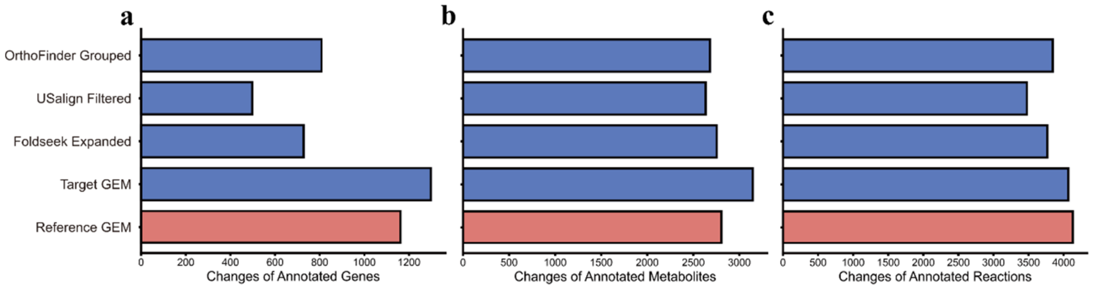


**d**

Distribution of reactions for GEMs constructed by AlphaGEM

| Strain | Total Reactions | Transport Reactions | Reactions with Isoenzymes | Reactions with Complex |
|---|---|---|---|---|
| *C. albicans* | 4066 | 1060 | 947 | 171 |
| *S. pombe* | 3694 | 1032 | 828 | 94 |

Supplemental Figure 4 GEM reconstruction for *C. albicans* using AlphaGEM.

Changes in the number of annotated genes (a), metabolites (b), and reactions (c) during the stepwise construction of the *C. albicans* GEM: OrthoFinder grouped; US-align Filtered; Foldseek expanded; final output GEM, along with the comparison with reference GEM: Yeast9. Distribution of different type of reactions in the constructed *C. albicans* and *S. pombe* models (d).

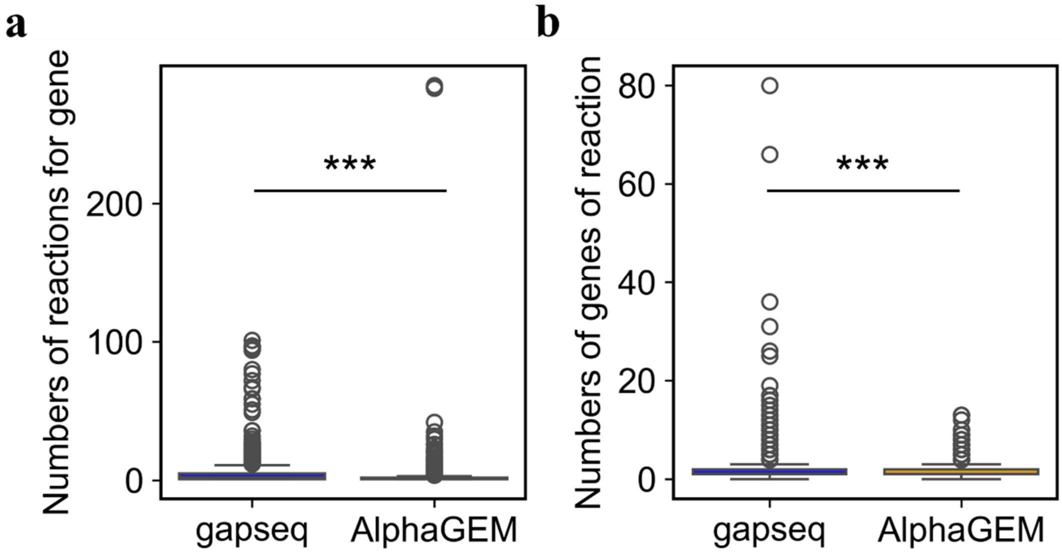


Supplemental Figure 5 Comparison of *K. pneumoniae* GEMs constructed by gapseq and AlphaGEM, respectively.

Distribution in the number of reactions catalyzed by one gene in GEMs built by gapseq and AlphaGEM, respectively, GEM built by gapseq has more extreme values (a). Distribution in the number of genes (including isozymes and enzyme complexes) catalyzing one reactions in GEMs built by gapseq and AlphaGEM, respectively, GEM built by gapseq has more extreme values (b). (***: *P* value < 0.001)

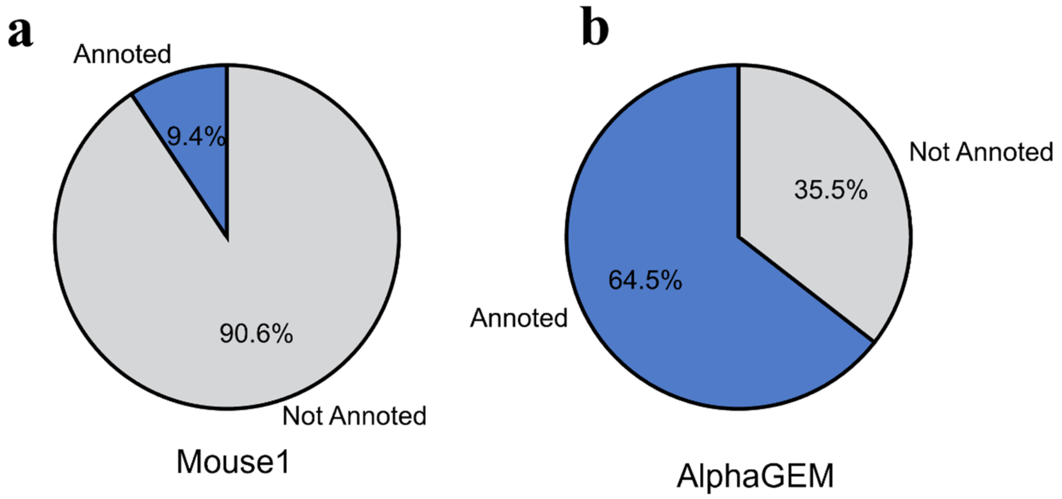


Supplemental Figure 6 Comparison of differential gene-protein-reactions in Mouse1 and GEM reconstructed by AlphaGEM.

Proportion of proteins annotated for different reactions in the Mouse1 model compared to AlphaGEM(a), in the AlphaGEM model (b). *Analysis performed by integrating multiple AI tools with a higher threshold (0.5) to identify differential reactions between models*

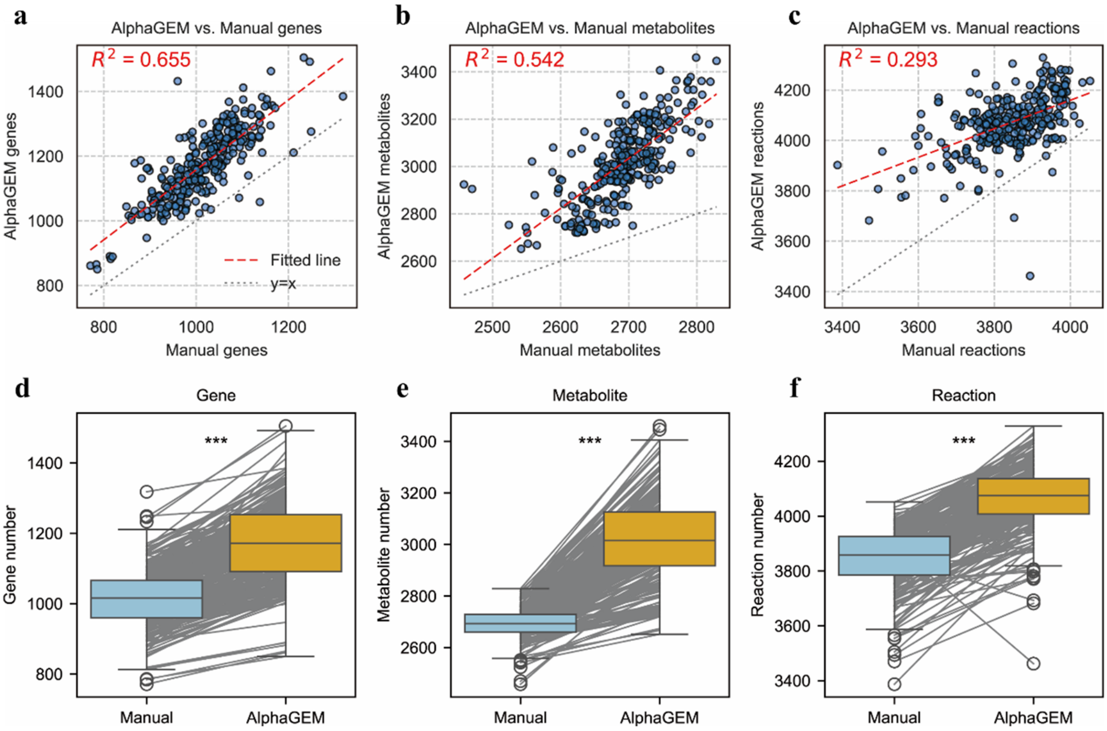


Supplemental Figure 7 Comparison and linear regression analysis between GEMs built by AlphaGEM and manual curation for 332 yeast species.

Correlation analysis in gene counts (a), metabolite counts (b), reaction counts (c) between GEMs built by AlphaGEM and manual curation. Comparison of gene counts (d), metabolite counts (e), reaction counts (f) between GEMs built by AlphaGEM and manual curation. (***: *P* value < 0.001)

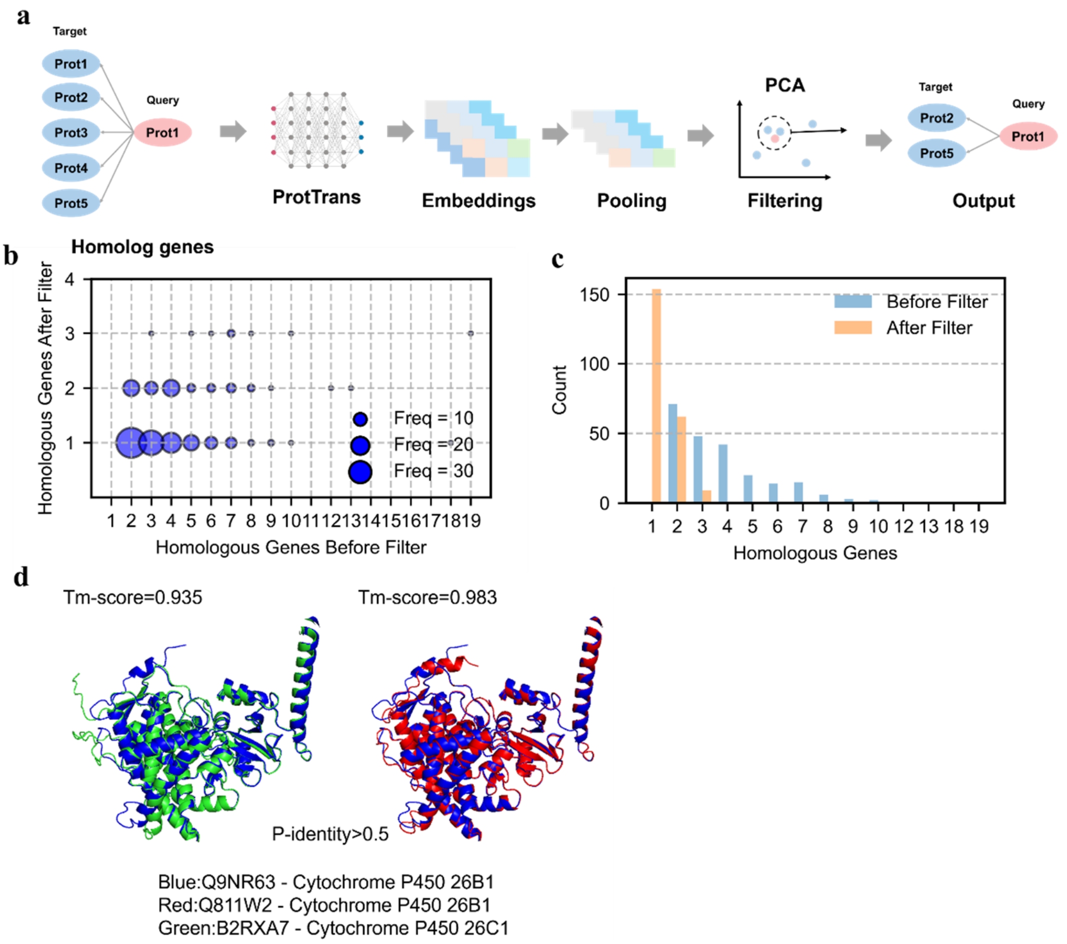


Supplemental Figure 8 Homologous proteins could be refined using ProtTrans.

ProtTrans-generated embeddings were used for proteins with multiple homologous proteins, followed by filtering based on a pre-selected distance threshold in PCA-reduced clustering space (a). Distribution of different changes in the number of homologous proteins per protein after filtering (b). Compare of global distribution in the number of homologous proteins per protein before and after filtering (c). This approach successfully eliminated erroneous homologous protein assignments derived from structural alignment during GEMs reconstruction (d).
